# Risk-reward trade-off in motility endurance generates dichotomy in search strategies among copiotrophic marine bacteria

**DOI:** 10.1101/2025.02.26.640445

**Authors:** Johannes M. Keegstra, Zachary C. Landry, Sophie T. Zweifel, Benjamin R.K. Roller, Clara Martínez-Pérez, Estelle E. Clerc, Martin Ackermann, Roman Stocker

**Affiliations:** Institute for Environmental Engineering, ETH Zurich, 8093 Zurich, Switzerland; Swiss Federal Institute of Aquatic Science and Technology (EAWAG), 8600 Duebendorf, Switzerland; Division of Microbial Ecology, University of Vienna, 1030 Vienna, Austria; Department of Environmental Systems Science, ETH Zurich, 8006 Zurich, Switzerland; School of Architecture, Civil and Environmental Engineering, Ecole Polytechnique Fédérale de Lausanne (EPFL), 1015 Lausanne, Switzerland

## Abstract

Coptiotrophic marine bacteria contribute significantly to carbon storage in the ocean by remineralizing organic carbon present in nutrient-rich hotspots amidst oligotrophic waters. Motility is both highly beneficial and costly in such environments, presenting copiotrophs with a risk–reward trade-off in search behavior. Here we studied the motility endurance of 26 marine isolates using video microscopy and cell tracking over two days of carbon starvation. We found that this cost-benefit trade-off results in a distinct dichotomy among marine bacteria: risk-averse copiotrophs ceased motility within hours (‘limostatic’), whereas risk-prone copiotrophs converted 10% of their biomass per day into energy to retain motility for the two days of observation (‘limokinetic’). We identified a genomic component of this dichotomy, sufficiently robust to predict the response of additional species with 83% accuracy and the prevalence of both strategies in the ocean. This dichotomy can facilitate the incorporation of the bacterial contribution in ocean carbon cycle models.

## Introduction

There is a profound dichotomy in ecological strategies among marine bacteria between oligotrophic and copiotrophic bacteria [1], representing a fundamental simplifying principle in our understanding of microbial ecology in the ocean. This dichotomy is associated with a suite of ecological and behavioral adaptations: oligotrophic bacteria are frequently smaller, non-motile, and mainly forage through molecular diffusion [2], and can thus more readily survive in the more oligotrophic regions of the ocean. Copiotrophic bacteria are larger, often motile, and have a high level of signaling and regulation to respond to changes in the environment [1], and are thus better adapted to feast–famine cycles driven by encounters with resource-rich hotspots [3, 4]. Between hotspots, copiotrophs experience strong growth limitation due to nutrient or energy starvation [5, 6], which may last for several days when the concentration of hotspots is low, such as in the open ocean [7].

Flagellar motility [8] can be highly beneficial for navigating heterogeneous environments [4], but is associated with a high demand on cellular resources [9, 10, 11]. Bacteria require an estimated 10^4^−10^5^ ATP/s for swimming in the absence of growth [9]: this energetic cost is comparable to or greater than the total cellular maintenance energy flux of 10^4^ ATP/s [12, 13], making flagellar motility the most expensive cellular trait during starvation. The evidence for motility during nutrient limitation has been mixed. It has been shown that some bacteria increase their investment in motility with decreasing growth rate due to nutrient limitation [14], and some species have been reported to retain motile behavior during starvation [15, 16]. However, most experiments to date show that starvation hampers motility [17, 18, 19, 20, 21, 22, 23]. Despite the high energetic requirements, motility potentially brings great rewards in the marine environment, by enhancing the encounter rate with localized nutrient hotspots such as phytoplankton cells [24] or organic matter particles [25] by 10^2^- to 10^3^-fold [26, 7, 27, 28]. These hotspots provide marine bacteria with rich nutrient resources: a marine particle can provide enough carbon for 10^3^ to 10^4^ new bacteria, meaning a successful colonization may lead to a manyfold increase in biomass [25]. This high potential search reward, combined with the risk of wasting limited cellular resources, makes bacterial motility under starvation subject to a risk–reward trade-off, and raises the question of which strategy is adopted by marine bacteria.

Here, we report on the motility behavior upon carbon starvation for 26 strains of 18 species of copiotrophic marine bacteria. We did not find a continuum of endurance timescales, but rather that a split in behavior exists between species that cease motility within a few hours and species that retain motility for multiple days, revealing an ecological dichotomy among motile copiotrophic bacteria. This dichotomy reflects a different risk assessment of starvation by different bacteria: risk-averse foragers cease motility to conserve resources until conditions improve, whereas risk-prone foragers retain motility to enhance their chance of large search rewards.

## Results

### Behavioral split in motility endurance upon carbon starvation

We measured the motility response of different marine bacteria to carbon starvation. Carbon starvation was imposed experimentally by growing the cells to exponential phase in carbon-replete Marine Broth (MB) then washing and placing the cells in carbon-deplete starvation medium. This procedure models for example the rapid loss of access to nutrients that cells experience when leaving a nutrient hotspot (Fig. 1A). We sampled cells immediately before washing and then at 1 h and approximately 3, 7, 22, 30 and 46 h after the onset of starvation. For every time point, we used video microscopy and cell tracking to quantify the cellular velocity (the velocity averaged over the cell’s trajectory) of approximately 300 cells. Our measurements reveal a striking divergence in the motility response to starvation, even among closely related species. As an example between closely related species (see the phylogenetic tree in Fig. S1), in carbon-replete medium, *Vibrio splendidus* FF-500 and *Vibrio anguillarum* 12B09 (previously known as *Vibrio ordalli* [29]) were both highly motile, with population-averaged velocities of 29±18 µm/s and 41±21 µm/s, respectively (Fig. 1B, Supplementary Videos S1 and S2). However, their motility upon entering carbon starvation was strikingly different. Within 1 h of starvation, the velocity of *V. splendidus* FF-500 diminished to 4±4 µm/s, whereas the velocity of *V. anguillarium* 12B09 during the two days of starvation remained high, with an average of 31±21 µm/s (Fig. 1B, Supplementary Videos S3 and S4). Experiments with an additional pair of strains from the same two species showed similar results (Fig. S2A). These observations show that bacterial species can have strongly divergent motility responses upon carbon starvation.

**Figure 1:**
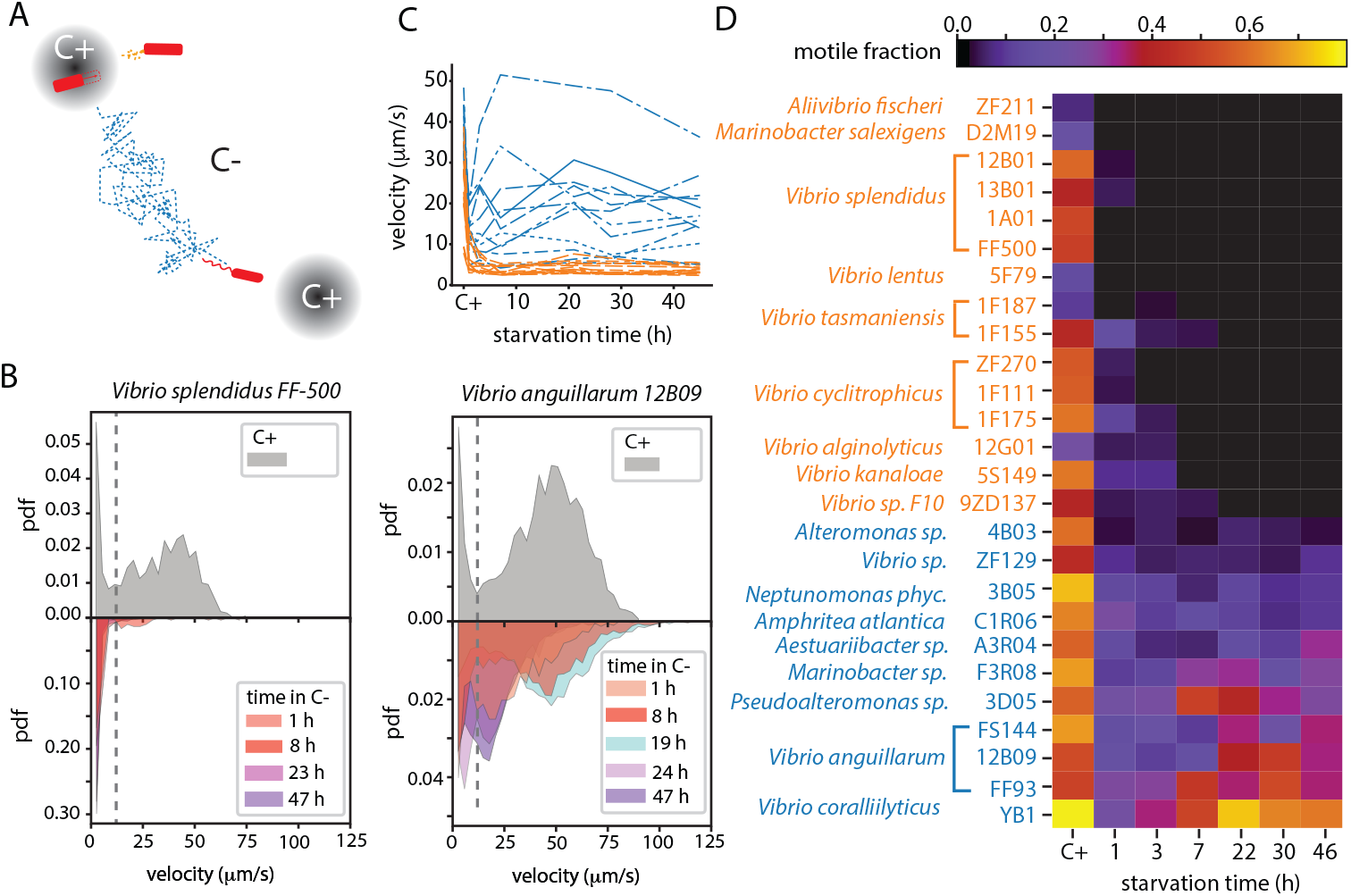
Marine bacteria exhibit a dichotomy in their motility response to carbon starvation. (*A*) Marine bacteria often experience carbon starvation (C-) during the time between encounters with carbon-replete hotspots that support growth (C+, gray circles). During starvation, bacteria may opt to cease motility (orange) to conserve resources or to sustain motility (blue) to increase chances of encountering a hotspot (*B*) Distribution of cellular velocities in *V. splendidus* FF-500 (left) and *V. anguillarum* 12B09 (right) prior to starvation (C+; top) and at different times during carbon starvation (1 h to 47 h; bottom). Dashed gray lines mark the velocity of 12 µm/s, used to differentiate motile from non-motile cells (Fig. S2B). ‘pdf’: probability density function. (*C*) Average cellular velocity of the population as a function of starvation time for 26 marine strains. 15 strains show a rapid decrease of velocity (orange), to on average 5±2 µm/s, whereas 11 strains retain a high velocity (blue), with an average of 18±9 µm/s. (*D*) Fraction of motile cells (as given by the colorbar at top) for the 26 strains as a function of starvation time. ‘C+’ denotes the condition prior to starvation. A total of 547 video microscopy experiments were performed; the number of experiments for each condition (typically 2–5 per strain) are given in Fig. S1.

We performed these carbon starvation experiments to measure the motility of 26 strains from 18 species belonging to the Gammaproteobacteria class. All strains exhibited a run-reverse-flick motility pattern (Supplementary Videos S5-S10), a hallmark of single-flagellated bacteria [30, 31]. For each strain, we computed the population-averaged velocity as a function of starvation time (Fig. 1C). We then used the cellular velocity distribution to distinguish actively swimming cells from those moving only by Brownian motion, identifying cells with a velocity greater than 12 µm/s as motile (dashed lines in Fig. 1B and Fig. S2B). The motile fraction for a single time point is defined as the frame-averaged ratio of motile to total cells (Materials and Methods). Following carbon starvation, the fraction of motile cells revealed a clear dichotomy: for some strains the motile fraction decreased rapidly to near zero, whereas for other strains it remained considerably above zero throughout starvation (Fig. 4D, Supplementary Videos S5-10).

To have an objective criterion to determine which strains retained and which strains ceased motility, we computed the kernel density estimate (KDE) of the log-transformed motile fraction, averaged for all starvation times exceeding 1 h, for each strain. The KDE exhibits a bimodal distribution with a minimum at a motile fraction of 0.033, providing a clear separation into two classes (Fig. S2D). We used this criterion to separate the motility response of each strain into two classes: 15 out of 26 strains had a motile fraction below this threshold upon starvation (on average 0.01 ± 0.01; unless noted otherwise computed as the average ± one standard deviation (S.D.) of the average value per strain), whereas the remaining 11 out of 26 strains retained a higher motile fraction than this threshold (on average 0.23 ± 0.16) (Fig. 1D). We propose to call the motility-retaining response ‘limokinetic’ (from the Greek λιµóσ meaning ‘starvation’) and the motility-renouncing response ‘limostatic’.

In a carbon-replete environment, the 26 strains exhibited no dichotomy in motility behavior and all had a motile fraction of at least 0.07 (on average 0.48±0.19) (Fig. 1D, Table S1). However, the limokinetic strains were on average more motile than the limostatic strains before the onset of starvation, having both a higher fraction of motile cells (0.61±0.11 vs. 0.39±0.20) and a higher average velocity of the motile fraction (43±7 vs. 33±8 µm/s) (Fig. S2E,F, Table S1).

To test the robustness of the observed dichotomy, we repeated experiments by using a different treatment to impose carbon starvation. Instead of washing the cells and transferring them to starvation medium, we measured the motility of cells during stationary phase in MB medium, during which nutrients are limiting. For six limokinetic and six limostatic strains, we compared the time-averaged motile fraction for each strain in the stationary phase to that obtained in the previous experiments in starvation medium, and found that the values were highly correlated (Fig. S3, Pearson’s *ρ* = 0.91). Furthermore, the classification into limokinetic and limostatic strains (based on the motile fraction criterion) was the same under the two treatments, with a single exception (*Alteromonas sp*. 4B03 was classified as limostatic in stationary phase and limokinetic in starvation medium). These results indicate that the dichotomy is robust to differences in the mode in which carbon starvation is imposed and is primarily a species-specific trait.

### Differential flagellar loss indicates commitment to non-motile and motile lifestyles

Loss of flagellar filaments during nutrient limitation has been reported for other bacteria [21, 23], prompting us to investigate the flagellation of limokinetic and limostatic strains during carbon starvation. We quantified the flagellation in five strains using scanning electron microscopy (SEM): two limostatic strains (*V. splendidus* 1A01 and *Vibrio cyclitrophicus* ZF270) and three limokinetic strains (*Alteromonas sp*. 4B03, *V. anguillarum* FS144 and *Vibrio coralliilyticus* YB1) (Fig. 2A). As for most other marine bacteria [32], the dominant mode of flagellation was a single polar flagellum (Fig. 2A): among the 1104 cells imaged by SEM, we observed 572 cells with 1 flagellum, on average 3.9±1.5 µm long, 528 with 0 flagella, and 4 with multiple flagella (3 of which were cells about to divide).

**Figure 2:**
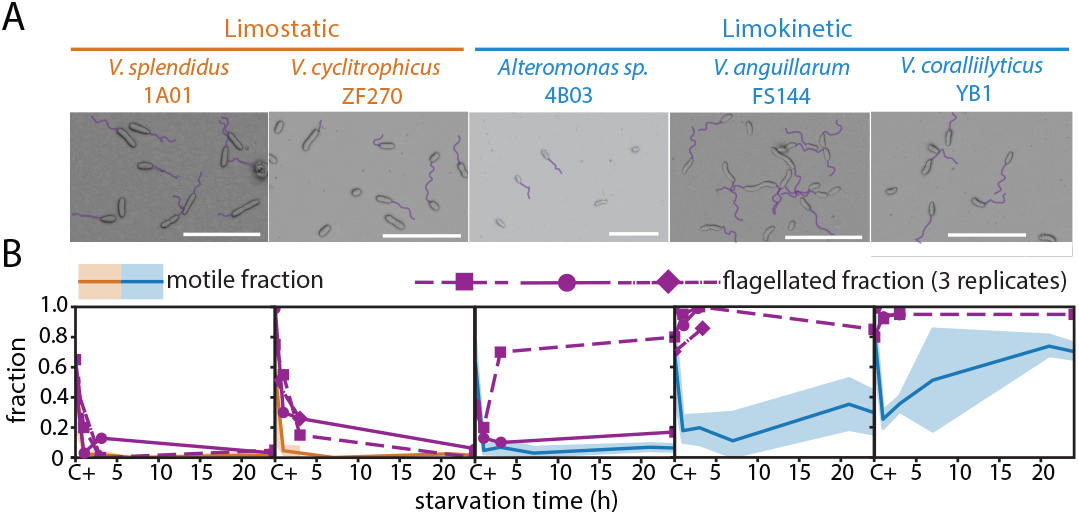
Limokinetic and limostatic strains differ in the prevalence of flagellation upon starvation. (A) Representative scanning electron microscopy images (SEM) from five strains, two limostatic and three limokinetic, from exponentially growing cultures (i.e., prior to the onset of starvation). Flagellar filaments are highlighted (purple). Scale bars = 10 µm. (B) Fraction of flagellated cells (purple lines) determined from SEM images as a function of starvation time, for two to three replicates per strain (symbols). ‘C+’ denotes conditions prior to starvation. The number of cells imaged by SEM per strain and time point was at least 34. For comparison, the fraction of motile cells for the same strains are shown (orange for limostatic, blue for limokinetic; data from Fig. 1D) along with 95% confidence intervals (CI; shaded areas).

The flagellation during starvation revealed a strong difference between limostatic and limokinetic strains. During exponential growth in a carbon-replete environment the flagellation was similar in the two classes, with an average flagellated fraction of 0.65±0.11 and 0.69±0.26 for limostatic and limokinetic strains, respectively. After 24 h starvation, the fraction of flagellated cells was only 0.04 ±0.01 in limostatic strains, whereas in limokinetic strains it was 0.75±0.20 (Fig. 2B). The average filament length did not show a difference between the two classes, nor was there a significant difference in filament length between starving or growing conditions (Fig. S4), indicating that during flagellar loss the filaments are lost in their entirety. Overall, this shows that during starvation limostatic strains lose flagellar filaments, whereas limokinetic strains retain them.

Comparing the fraction of flagellated cells with the fraction of motile cells suggests that bacteria can control motility independently from flagellation. For all strains the fraction of flagellated cells was higher than the fraction of motile cells (Fig. 2B). The difference is especially strong for the limokinetic strains *V. anguillarum* FS144 and *V. coralliilyticus* YB1, where the flagellated fraction was respectively 3.8 and 1.9 times larger than the motile fraction (FS144: 0.92±0.05 vs. 0.24±0.03; YB1: 0.51±0.06 vs. 0.95 ±0.01; mean ±one standard error of the mean (S.E.M.)) (Fig. 2B). This suggests that marine bacteria have the ability to pause motility, by temporarily stopping flagellar rotation, without flagellar loss.

We performed further experiments to exclude the possibility that the observed flagellar loss was due to shear stress [33, 34] induced by washing during the transfer to starvation medium. In control experiments where cells were washed using the same procedure yet with carbon-replete medium in the place of starvation medium, we observed instead an increase in the motile fraction (Fig. S4), demonstrating that the washing protocol (and associated shear stress) does not cause flagellar loss. This indicates that flagellar loss in our starvation experiments was a specific response to nutrient depletion, not to mechanical stress.

Flagellar loss prevents bacteria from rapidly responding when conditions improve, as flagellar synthesis is slow: even a relatively short flagellar filament of 1.5 µm-long requires at least 30 min to be synthesized [35, 36, 33]. To quantify the response delay, we starved three limostatic and eight limokinetic strains for 24 h, then exposed them to carbon-replete MB medium, to mimic the encounter with a nutrient hotspot. We quantified motility at 10, 30 and 60 min after this nutrient addition. For the limostatic strains, the fraction of motile cells remained low for the first 10 min and recovered only after 30 to 60 min. By contrast, five out of eight limokinetic strains exhibited an increase in motile fraction and swimming velocity 10 min after nutrient addition (Fig. S5). These observations indicate that limostatic strains not only stop swimming, but commit to a non-motile lifestyle, and as a result experience a significant delay (tens of minutes) to adapt back to carbon-replete conditions, in contrast to limokinetic strains that retain motility throughout.

### Limokinetic strains convert biomass into energy to fuel motility

During starvation, the synthesis of new motility machinery diminishes and the dominant cost of motility is the operation of the flagellar motor for propulsion [9]. The average power spent on motility per cell can be estimated as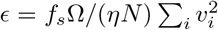, with *v*_*i*_ the average swimming velocity of motile cell *i, N* the number of motile cells, *f*_*s*_ the fraction of motile cells, *η* the efficiency of the flagellum (2% [37]) and Ω the resistance coefficient of the bacterium including its flagellum (Ω = 4.1·10^*−*8^ Nsm^*−*1^ [38]). We used the motile fraction and the swimming velocities (Figs. 1D, S7, Table S1) to compute the power spent on motility per cell, for each strain as a function of time (Fig. 3A). The energy expenditure of limokinetic strains on average decreased more than three-fold during starvation (from 4.1±0.4 · 10^4^ ATP/s before starvation to 1.2±0.4·10^4^ ATP/s during starvation, mean ±S.E.M.), assuming a conversion factor of 8· 10^*−*20^ J/ATP [9]). Even with this decrease, the energy required for motility was similar to the typical maintenance energy during starvation, estimated at 1·10^4^ ATP/s per cell [13, 12]. Hence, by remaining motile, limokinetic strains at least double their energy requirements during carbon starvation compared to limostatic strains.

**Figure 3:**
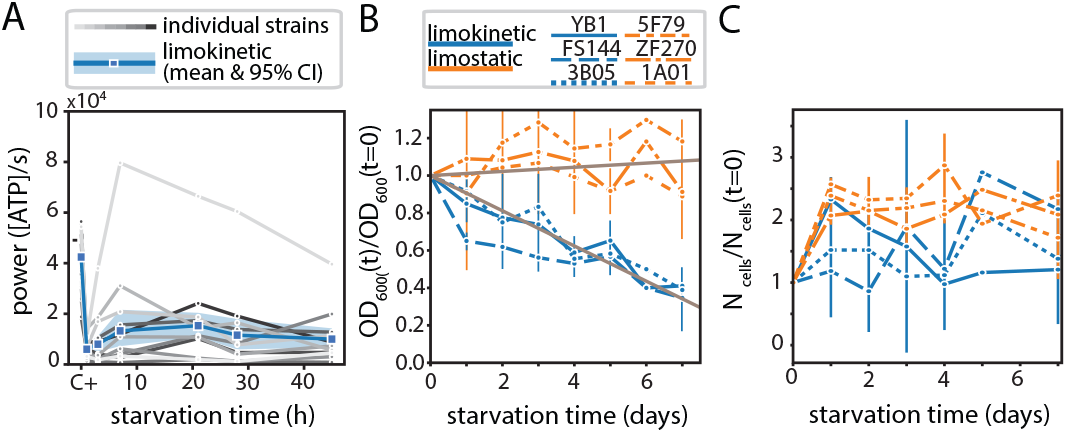
Limokinetic bacteria convert biomass to energy to fuel motility. (*A*) Estimated motility power requirement per cell for each of the 11 limokinetic strains (gray lines) and average over the 11 limokinetic strains (blue, 95% CI shown as shaded area), as a function of starvation time, for two days of starvation. (*B*) Optical density as a function of starvation time, normalised by the optical density at the onset of starvation (*t* = 0), for three limostatic (orange) and three limokinetic (blue) strains, for 7 days of starvation. Shown are the average and standard deviation (vertical lines) of three experiments. Different strains are denoted by different line types. Grey lines indicate linear fits to the change in optical density of limokinetic and limostatic strains, with slopes of respectively -0.094 d^*−*1^ and +0.011 d^*−*1^ (see main text). (*C*) The number of cells *N*_cells_ after prolonged starvation, normalized by the number of cells at the onset of starvation (*t* = 0), for three limostatic (orange) and three limokinetic (blue) strains, for 7 days of starvation. Strains and error bars as in panel B.

We hypothesised that cells use internal energy sources to fuel motility, sacrificing a part of their biomass to generate energy [39]. This hypothesis is supported by estimates of the biomass loss that would be required to fuel motility. A power requirement for motility of 1.2· 10^4^ ATP/s, corresponding to a motile fraction of ∼0.3 and a swimming velocity of ∼40 µm/s, would require a biomass loss of approximately 10 fg worth of glucose per day (glucose yields ∼30 ATP per molecule). Such a biomass loss would for example be insurmountable for marine oligotrophs, with typical cell mass of 10 fg [40]. For copiotrophs, instead, it represents a daily loss of only about 5% of their typical biomass of 200 fg [40]: this conversion would thus allow copiotrophic bacteria to retain motility for multiple days.

To test the hypothesis that limokinetic bacteria convert biomass into energy for motility, we used optical density as a metric for biomass [41] and measured the optical density of three limokinetic and three limostatic strains during a seven-day starvation experiment. The three limokinetic strains lost on average 62% ± 3% of their biomass. In contrast, the biomass of the limostatic strains remained approximately constant (99%±16%) (Fig. 3C). A linear fit of OD_*n*_ = 1−*γt* over all individual measurements on limokinetic strains yielded a good fit (*R*^2^ = 0.88), with a biomass decay rate rate of *γ* = 0.094 d^*−*1^ (95% CI: [0.085,0.103]). The same fit yielded *γ* = -0.011 d^*−*1^ (95% CI: [-0.025,0.003], *R*^2^ = 0.04) for the limostatic strains. All three limokinetic strains were motile over the seven-day period (motile fraction of 0.22±0.19, average swimming velocity 42± 7 µm/s, Fig. S8A), indicating that motility endurance was associated with a biomass loss of 9.4% per day and suggesting that limokinetic strains sacrifice a part of their biomass to fuel motility.

We confirmed that the biomass decrease is due to a conversion of biomass to energy, rather than a decrease in the number of cells. Flow cytometry measurements of the cell number during the seven-day starvation experiment revealed that the number of cells increased or remained constant compared to the onset of starvation (Fig. 3C). After seven days of starvation, the increase in cell number (over the initial cell number) was 1.59±0.51 for limokinetic strains and 2.06±0.33 for limostatic strains. Alternative estimates based on colony counts and the number of cell tracks confirmed that the number of cells did not decrease during starvation (Fig. S8). The increase in cell number was likely due to reductive divisions, a well-known starvation response where the population biomass is redistributed over more, but smaller, cells [5, 23]. Our results reveal a starvation response that additionally reduces the cellular biomass in limokinetic strains, due to a conversion of biomass to energy to fuel motility.

Additional experiments allowed us to exclude three alternative energy sources for motility. First, we considered the recycling of necromass [42, 43]. Live/dead staining showed that the fraction of dead cells was comparable between the two classes (0.07±0.06 and 0.11±0.07 for limostatic and limokinetic strains, respectively, when averaged over the week of starvation (Supplementary text, Fig. S8E). Given the small difference in death rates, and considering that necromass recycling is typically inefficient (10– 20%) [43], this means necromass recycling does not represent a large energy source for motility in our starvation experiments. Second, we found that the energy source is not photonic in nature, as limokinetic strains lack rhodopsin genes and remained motile when starved in the dark (Supplementary Text, Fig. S9A). Third, we excluded the effect of any residual nutrients in the starvation buffer, by showing that there was no negative dependence of the motile fraction on cell concentration (Supplementary Text, Fig. S9B).

To understand the nature of the biomass that is converted into energy, we investigated whether limokinetic strains accumulate energy storage compounds before starvation. Using four limokinetic and three limostatic strains in carbon-replete media, we assayed the presence of polyphosphate (polyP) using DAPI staining and polyhydrobutyrate (PHB) using Bodipy staining. Both storage compounds have been suggested to act as energy sources for motility [44, 45]. We observed no formation of polyP granules (Fig. 4A), except in 4% of cells in limokinetic strain *Alteromonas* sp. 4B03 (Fig. S10). In contrast, two out of three limokinetic strains produced considerable amounts of PHB, whereas limostatic strains produced either no PHB or lower amounts (Fig. 4B,C). This suggests a potential role of storage compounds as an additional yet not exclusive energy source of motility in limokinetic strains during starvation. Another bacterial storage compound is glycogen, for which many limokinetic and limostatic strains have synthesis genes (Table S2) but it is unclear if glycogen can act as an energy source over extended times [46, 47]. Finally, bacteria may catabolize biomass that does not have a specific storage function, such as lipids or ribosomes, for energy [5].

**Figure 4:**
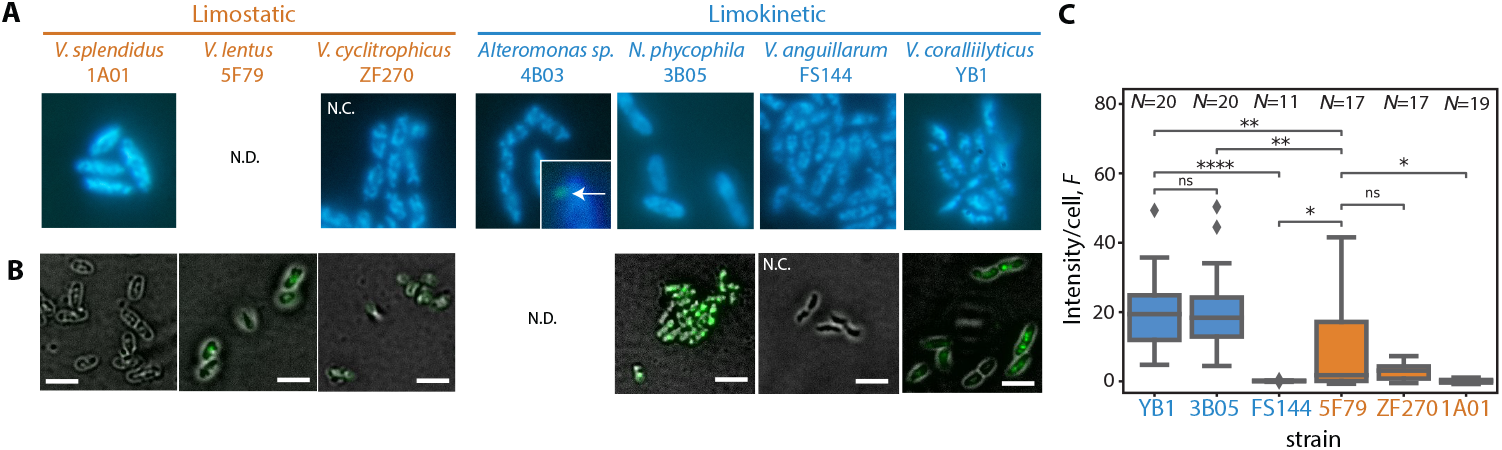
Accumulation of energy storage compounds in carbon-replete medium. (*A*) Absence of polyphosphate (polyP) storage during exponential growth in rich medium in two limostatic and four limokinetic strains, measured by DAPI staining. PolyP granules are visible in green, DNA in blue. None of the inspected cells showed polyP granules, except for 2 out of 54 cells of strain 4B03 (one cell in inset, granule indicated by white arrow). N.C.: negative control (strain lacks storage compound synthesis genes, see Table S2). N.D.: Not determined.(*B*) Polyhydrobutyrate (PHB) storage in three limokinetic and three limostatic strains during exponential growth in rich medium, measured by Bodipy staining. Scale bars = 3 µm. (*C*) Fluorescence intensity per cell from staining of PHB with Bodipy, for each strain. Box plots represent the values of the first, second (median) and third quartiles. Whiskers indicate minimum and maximum values of the distribution, limited by 1.5 times the difference between the first and the third quartile. Diamonds indicate outliers. Sample sizes indicate the number (*N*) of individual cells measured. Significance based on bidirectional M.W.U: * *p <* 0.05; ** *p <* 0.01; **** *p <* 0.0001; ns = not significant.

### The genomic basis of the limokinetic and limostatic lifestyles

We investigated the genomic basis of the difference between limokinetic and limostatic behaviors using assembled genomes of all strains, in order to identify further differences between limokinetic and limostatic strategies and to predict the motility behavior under starvation in other bacterial species. We constructed a Bayesian classifier, selecting genetic features that are associated with limokinetic behavior through recursive feature elimination (RFE; Materials and Methods). The classifier was able to separate the two behavioral classes with good accuracy (85%, Figs. 5A, S11), defined as the fraction of correctly predicted strains.

**Figure 5:**
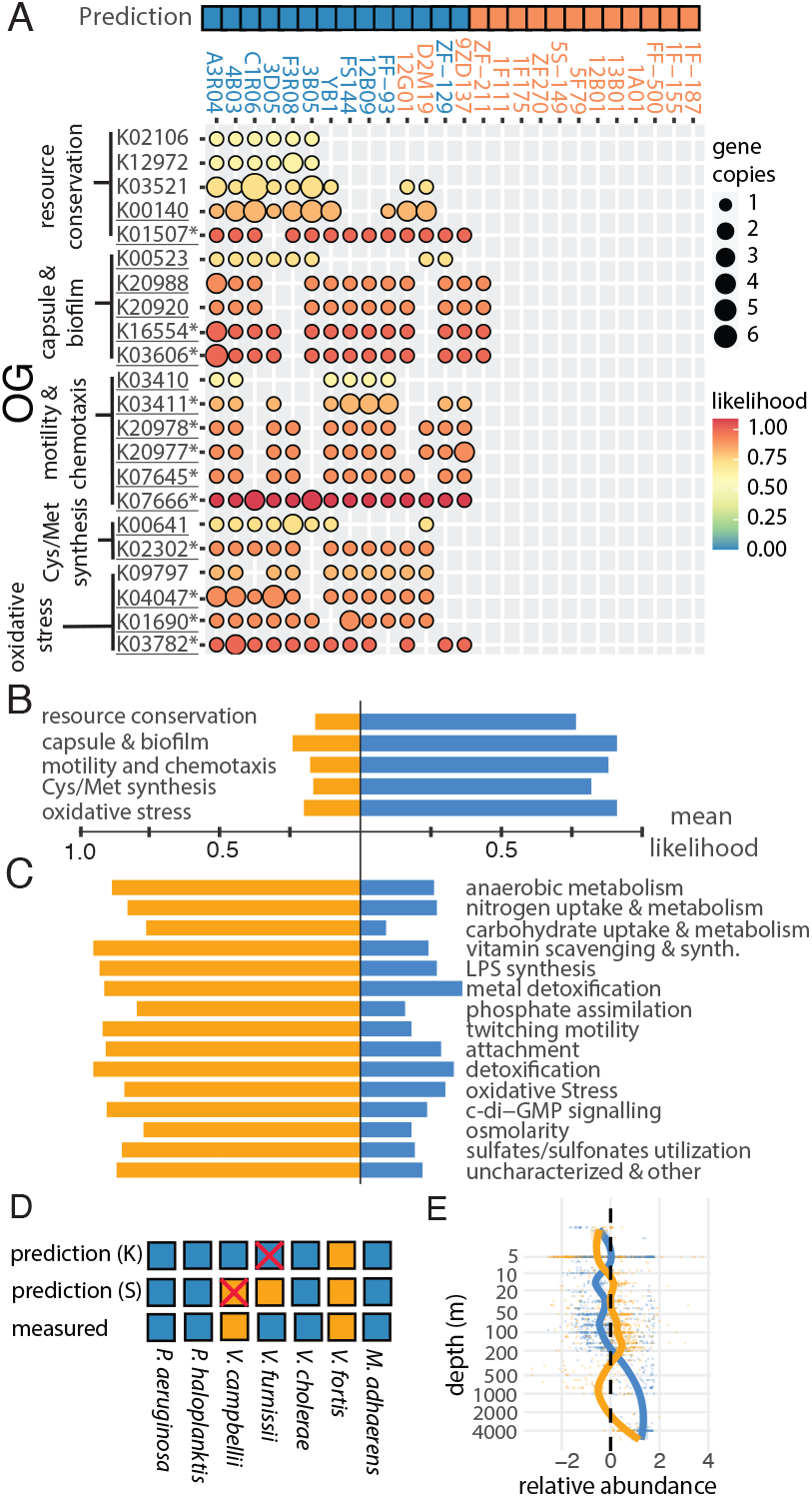
Genomic basis of the limokinetic–limostatic dichotomy. (*A*) Bayesian classifier for the prediction of limokinetic behavior. (top) Classifier prediction of the limokinetic (blue) and limostatic (orange) behavior for all strains. Strain names are color-coded according to their experimentally determined classification (Fig. 1). (bottom) Prevalence of orthogonal groups (OG) associated with a limokinetic response for both limokinetic and limostatic strains, as obtained by RFE (Materials and Methods) and clustered into five functional categories (Supplementary Text). Circles indicate the gene copy number of each OG (size) and the probability of association with the limokinetic response (color). Underlined OGs indicate significance corrected for phylogeny of *p <* 0.10 and marked by * indicate *p <* 0.05 (Table S3, Materials and Methods) (*B*) Mean likelihood averaged over all OGs in each functional category, for both limokinetic (blue) and limostatic (orange) strains, as predicted by the classifier for the limokinetic response (panel A). (*C*) As in B, but for a classifier based on genes associated with the limostatic response (classifier features in Fig. S13). (*D*) Prediction of the motility response to starvation based on the genomic classifiers for seven strains not included in classifier training (squares), for both the limokinetic (‘Pk’) and limostatic (‘Ps’) classifiers. (*E*) Predicted relative abundance (z-score) of limokinetic (blue) and limostatic (orange) taxa as a function of depth, computed by applying the classifiers to 1038 field samples from the Ocean Microbiomics Database [49]. For limostatic taxa, the z-score of individual samples (dots) is computed as the abundance of limostatic taxa after subtraction of the depth-averaged abundance and normalization to one standard deviation. The same procedure is applied for limokinetic taxa. For both strategies, a moving-average (solid lines) computed with a Loess filter using a window of 2/3 of the data is also shown. The relative abundances of limokinetic and limostatic taxa are anti-correlated with depth (Pearson’s *ρ* = −0.79).

The classifier relies on a set of 22 orthologous groups (OG), or genes with conserved function, associated with a limokinetic response to carbon starvation (Fig. 5A). We grouped the genes into five functional categories based on their annotated function (Supplementary Text). Genes selected by RFE include mechanisms for resource conservation (*N*_O_ = 5 genes, average likelihood L = 0.76), capsule and biofilm formation (*N*_O_ = 5, L = 0.91), regulatory elements of motility and chemotaxis (*N*_O_ = 6, L = 0.88), cysteine/methionine synthesis (*N*_O_ = 3, L= 0.82) and oxidative stress response (*N*_O_ = 3, L= 0.91) (Fig. 5B, Supplementary Text).

Oxidative stress is abundant in the marine environment [48] and oxidative stress defense genes have a high associated likelihood in the classifier. Therefore, we tested if the limokinetic and limostatic strains exhibited different sensitivity to oxidative stress. We found only a mild (∼25%), non-significant (*t*-test, *p* = 0.23) increase in sensitivity for limostatic strains, measured as the increase in lag time due to oxidative stress (Fig. S12, Supplementary Text). It is possible, however, that the cumulative effects of oxidative stress may only have an impact in cells subjected to long-term energy and carbon starvation, where the ability to repair or synthesize new proteins is compromised.

We also trained the classifier in the inverse direction, selecting for genes that are associated with a limostatic response to starvation. The accuracy of this limostatic classifier (85%) is similar to the limokinetic classifier. The 124 genes selected by RFE are implicated in a number of cellular functions (Fig. 5C), including anaerobic metabolism (*N*_O_ = 23, L = 0.87), and the uptake and metabolism of amino acids (*N*_O_ = 15, L = 0.86) and carbohydrates (*N*_O_ = 14, L = 0.78). Furthermore, there are genes involved in type IV pili attachment (*N*_O_= 4, L= 0.78) and twitching motility (*N*_O_ = 6, L= 0.92) (Figs. 5C, S12, Supplementary Text). Together, this could indicate an increased metabolic versatility and surface specialization of limostatic compared to limokinetic strains.

Because there is a taxonomic signature in our data (26% of the Vibrionaceae family are limokinetic, compared to 86% of strains from other families; Fig. S1), we considered the effect of phylogeny on feature selection using *post-hoc* phylogenetic logistic and linear regression analyses of classification outcome against OG presence (Materials and Methods). Of the 22 and 128 OGs in the limokinetic and limostatic classifiers, respectively, 12 and 56 OG’s have relationships with classification outcome cannot be explained by phylogeny alone with high certainty (p *<* 0.05) and 6 and 14 with moderate certainty (p *<* 0.10). (Table S3, Supplementary Text). Furthermore, our classifiers performed better than a taxonomic classifier (*p* = 0.07) and were able to accurately predict variation within the Vibrionaceae (Supplementary Text). We therefore expect our classifier to be applicable to other motile marine bacteria.

We used the classifiers to predict the motility response to starvation of seven additional marine strains not included in the training of the classifiers. The prediction of the limokinetic classifier was correct for all tested strains except *Vibrio campbellii*, and the limostatic classifier predicted all marine strains correctly except *Vibrio furnissii* (Fig. 5D, accuracy 86% for both). Interestingly, the classifier predictions for the enteric species *Escherichia coli* and *Salmonella typhimurium* were ambiguous, in that these strains were predicted to be both limokinetic and limostatic. In experiments, their motile fraction decreased much slower (11-14 h) compared to the limostatic marine strains (2.4h, *p <* 1·10^7^,t-test) (Fig. S14B). This contrast with enteric bacteria indicates that the dichotomy we have described is a feature of marine bacterial communities: the extent to which it may occur in other microbiomes will require dedicated investigation.

Finally, we used the Global Ocean Microbiomes dataset [49] in combination with our classifiers to predict the prevalence of the two strategies among assembled metagenomes in the ocean (Materials and Methods), as the classifier was originally trained and tested using gammaproteobacterial taxa we limited our prediction to this group. Limokinetic taxa are predicted to dominate in 97.3 % of 1038 field samples (Fig. S15). This finding suggests that the limokinetic foraging strategy in the natural environment is more prevalent, in contrast to the findings from most laboratory-based studies predicting motility loss upon nutrient depletion [17, 18, 19, 20, 21, 22, 23]. However, certain environments may favor a limostatic strategy, as indicated by the fact that the samples with limostatic dominance all come from the euphotic zone (geometric mean depth of 66 m). The relative abundances of limokinetic and limostatic taxa as a function of depth are anti-correlated (Pearson’s *ρ* = −0.79, Fig. 5E), suggesting the presence of environmental variables that affect the abundance of both strategies. This could for example be due to the concentration of dissolved and/or particulate nutrients, yet more work is needed to determine the environmental drivers of the prevalence of one versus the other strategy.

## Discussion

Dichotomies are widely used as simplifying principles to help understand the daunting diversity of microbes in natural environments. Our results reveal an important dichotomy that separates motile copiotrophs into limokinetic and limostatic species. Both have the ability to swim, but show distinct behaviors in the face of carbon starvation: limostatic species rapidly cease swimming upon starvation, on a timescale of hours, whereas limokinetic species continue to swim, over a period of days. The consequence of the swimming endurance is the conversion of biomass to energy in limokinetic species (possibly enabled, at least in some cases, by storage compounds) versus the conservation of biomass in limostatic species. Our analysis of genomic signatures suggests that the limokinetic/limostatic dichotomy may be connected to other traits associated with oxidative stress defense, metabolism, uptake and surface-associated lifestyles. We have used these genomic signatures to construct classifiers to predict limokinetic or limostatic behavior in natural environments.

Traditionally, bacterial motility and chemotaxis have been understood as strategies to enhance foraging particularly in nutrient-poor environments [50, 51], yet recent work has shown the additional benefit of motility in nutrient-replete environments [52], mainly through enhanced colony expansion into uncolonized regions of the environment [53, 54]. Limostatic strains appear to use motility to disperse and colonize hotspots only during growth, when most of them are motile. This strategy may be especially effective under algal bloom conditions, when the number of hotspots is high (Supplementary Discussion) and so is the background level of dissolved organic matter, alleviating starvation. Limokinetic species likely also use motility for this purpose, but unlike limostatic species, we propose that they also use motility to actively search for hotspots in oligotrophic environments, even at the expense of sacrificing a sizeable fraction of their biomass to fuel motility. In such environments, limostatic copiotrophs will instead cease to be motile and conserve biomass until conditions improve again. Marine bacteria have been observed to survive starvation for periods of up to several months [55], suggesting that limostatic strains are specialists in overcoming large temporal distances, whereas limokinetic strains are specialists in overcoming larger spatial distances. While certain environments will favor one or the other phenotype, some of the limokinetic and limostatic strains studied here were isolated together (Table S1), suggesting that the two behaviors can co-exist.

Limokinetic behavior can be beneficial for marine bacteria to survive in the open ocean, where the concentration of particles is low and thus the search time for a new particle is high, up to multiple days or more [7]. All limokinetic strains were motile after 48 h (Fig. 1) and at least some can remain motile for one week (Fig. S8A), suggesting that they can successfully search for particles even in rather dilute environments. Furthermore, retaining motility for a long time is beneficial even if the average search time for a particle is longer still, because even a small fraction of cells encountering a particle that provides a very large nutrient return can allow a population to survive [7]. For example, if the average search time is 4 days, 10% of the motile population will encounter a particle in less than 9 hours (assuming exponentially distributed waiting times). Combined with the relatively fast division times of marine copiotrophs (*<*1 h in this study), these early arrivals are most important for the population dynamics on particles: a cell that arrives 12 h later than a (lucky) competing cell will have to overcome a setback of 2^12^ ≈4000 cells. Therefore, we believe motility during starvation aids limokinetic strains to survive even in very dilute environments, including ones in which particle concentrations are very low and thus search times very high.

This study has focused on the effect of starvation on motility alone, but what is the role of chemotaxis in the dichotomy? In the typical resource landscape of the ocean, hotspots are characterized by local gradients that marine bacteria must first find, before they can use chemotaxis to climb the gradient and home in on the hotspot: it is swimming per se (i.e., ‘random motility’), rather than chemotaxis, that provides the largest boost (100-to 1000-fold; [7]) in the encounter rate with such hotspots over non-motile bacteria. To increase encounters further, bacteria could either increase their swimming velocity, which is energetically costly, or suppress reorientation events (since this results in exploration of a larger volume [56], see Supplementary Discussion), which comes at no energetic cost. Our data reveal no such suppression: the average reorientation frequency of all limokinetic strains increased from 2.3±0.4 in nutrient-rich media to 3.3±0.8 after two days of starvation (*t*-test: *p* = 0.002, Fig. S7B, Table S1). Because reorientations are crucial for chemotaxis, a decrease in reorientation frequency would come at the cost of reduced responsiveness to short-lived chemical gradients [56, 57], which are typical in the marine environment [4, 58]. All 26 tested strains possess chemotaxis genes and both classes include species that are highly chemotactic, including for example *Vibrio anguillarum* [58] and *Vibrio alginolyticus* [59]. The fact that limokinetic strains do not suppress reorientations during starvation, despite the benefit this would provide in searching for hotspots, suggests that maintaining the ability to perform chemotaxis also during starvation is important. Furthermore, several limokinetic strains possess an alternative adaptation system (CheC, K03410 and CheD, K03411; Fig. 5A) that requires less resources and therefore could be beneficial for chemotaxis during starvation (Supplementary Text).

The flagellar loss in limostatic strains indicates that non-energetic reasons also influence the dichotomy in motility endurance. Upon starvation, limostatic strains lose flagella, instead of temporarily pausing motility, indicating a prolonged commitment to a non-motile lifestyle. Many species of bacteria are able to pause motility without losing flagella [60, 18, 61], likely through a ‘clutch-like’ mechanism [62, 63]. Our observation that during starvation the flagellated fraction of cells in limokinetic strains is higher than the motile fraction indicates that these strains are also capable of pausing motility. The ability of bacteria to control motility separately from flagellation suggests that bacteria do not need to eject flagella in order to halt motility to save energy, contrary to what has been proposed [21, 64]. There must thus be non-energetic reasons for the ejection of flagella under starvation, such as the avoidance of predation, which plays a significant role in oligotrophic environments [65]. Motility can affect predation by increasing encounter rates with predators [66, 67], but it is possible that even the mere presence of a flagellum could increase predation risk, for example by bacterivores [68] and phages [69, 70]. Therefore, it is possible that cells eject their flagella to decrease predation risk. This indicates that the dichotomy between limokinetic and limostatic behavior is shaped not only by energetics, but likely also predation pressure.

Our results highlight a dichotomy in bacterial motility behavior that results from a risk assessment between the anticipated biomass gain of motile behavior and the biomass loss due to conversion to energy and possibly predation. The dichotomy serves as a simplifying principle that can help predict the ecological and biogeochemical functions of marine microorganisms in the face of their astounding degree of diversity.

## Materials and Methods

### Bacterial cell culture and starvation protocol

Cells were inoculated from a frozen (−80 ^*°*^C) glycerol stock and grown overnight in 100% Marine Broth (BD Difco, Fischer Scientific) at 27 ^*°*^C on a rotary shaker (200 rpm). On the day of the experiment, cells were diluted 1/100 into half-strength Marine Broth with 50% Artificial Seawater (Instant Ocean, Aquarium Systems Inc., hereafter ‘ASW’). After 3.5–4 h, the cultures reached mid-exponential phase (OD 0.1–0.5) and were harvested by centrifugation (5000*g* for 6 min). Pelleted cells were resuspended in starvation medium, consisting of f/2 minimal medium without carbon (made by supplementing ASW with nitrogen, phosphorous, trace metals and vitamins (Provasoli-Guillard f/2 Media Kit, NCMA), with added 1 mM NH_4_Cl). This washing protocol was repeated three times, after which bacterial cells were diluted ten-fold compared to the original culture (leading to a cell concentration of ∼10^7^ cells/mL) and placed in a shaking incubator (175 rpm) at room temperature for the duration of the experiment. Bacteria were sampled from the medium immediately before washing the cells and at 1 (1), 2–4 (3), 5–9 (7), 19–24 (22), 28–32 (30), 43–48 (46) hours after the washing protocol started, where the number in brackets refers to the weighted average of each time window, rounded to 1 h, that was used for averaging over multiple experiments. Bacteria were observed within 15 minutes after sampling. The optical density of bacterial cultures was measured with a cuvette-based spectrometer (WPA Biowave Cell Density Meter, Biochrom Ltd.) on samples that were starved as described above but without the final dilution step (leading to an OD of 0.05–0.4, corresponding to ∼ 10^8^ cells/mL).

### Choice of bacterial strains

The strains used in experiments originate mainly from two principal collections. First, a collection of Vibrionaceae isolated off the Massachusetts coast [71], which has been extensively characterized for antagonistic interactions [72], colonization–dispersal behavior [73], and alginate degradation [74]. Second, a collection of coastal seawater isolates associated with chitin particles [75]. We also included *Vibrio coralliitycus* YB1, a highly motile strain isolated from corals [76]. Using publicly available genomes, we selected those likely to be motile, based on their number of motility and chemotaxis genes. Of the 107 available strains, we selected 36 strains to test for motility and growth, some of which were from the same species to encompass intra-species and inter-species variation, and all with both chemotaxis and motility genes. Of the 30 remaining strains (four strains did not grow in marine broth and two strains did not show motility during growth in marine broth), we randomly selected 26 strains to be used in this study.

The following species were selected to test classifier predictions (but were not used to train the classifier): *Vibrio fortis* KT626460, isolated from a healthy coral [77]; *Pseudoalteromonas haloplanktis* ATCC 700530 a model organism for chemotaxis studies in marine bacteria [78]; *Vibrio cholerae* C6706 was a gift from K. Drescher (U. Basel); *Vibrio campbellii* BB-120 was a gift from K. Jung (LMU, Munich); *Vibrio furnissii* was obtained through the German Collection of Microorganisms and Cell Cultures (DSMZ, 14383); *Marinobacter adhaerens* HP15 is a model organism for algae–bacteria interactions [79] and a gift from Matthias Ullrich (Jakobs University, Bremen). The HP15 strain contained a YFP-encoding plasmid but was grown and measured as for other non-fluorescent bacteria.

### Microscopy and cell tracking

Cell samples of 45 µL were placed in the center of a chamber (created by fixing a coverslip on a standard microscopy slide separated by silicone rubber of 1 mm thickness) and observed using phase-contrast microscopy (Nikon, JP) with a 20X (0.45 NA) air objective (S Plan Fluor ELWD, Nikon, JP). For very high cell densities and very low densities, 10X (0.30 NA) and 40X (0.60 NA) objectives were used, respectively. Videos with acquisition rate 25–30 frames per second were recorded using a CMOS camera (ORCA Flash 4.0, Hamamatsu) for 30 s, with a resolution of 2044 ×2048 pixels (0.326 µm/pixel for 20X). Cell tracking was performed using TrackPy (v 0.5.0, [80]) after removing the background from each image by subtracting the median image computed over the entire video. In the analysis, a maximum displacement per frame of 31 pixels (corresponding to a swimming velocity of 200 µm/s) and minimum separation between particles of 51 pixels were allowed. Trajectories shorter than 10 frames were removed from the analysis. Trajectories were then corrected for drift and cell positions were averaged over a time window of 5 frames in calculation of the velocity. The cellular velocity was defined as the velocity averaged over its trajectory and the population-averaged velocity is the mean cellular velocity of a population. Cells with a cellular velocity lower than 12 µm/s were classified as non-motile (Figs. 1A, S2B). A velocity of 12 µm/s corresponds to an approximate apparent displacement of 1 pixel between frames, due to diffusion and/or localization inaccuracy. For each frame *i*, the number of motile (*N*_*m,i*_) and nonmotile (*N*_*s,i*_) cells was determined. The motile fraction was then defined as (1*/T*) _*i*_ *N*_*m,i*_*/*(*N*_*s,i*_ + *N*_*m,i*_), where *T* is the total number of frames in the video. The average swimming velocity was defined as the average cellular velocity of all motile cells. Videos with a motile fraction lower than 0.075 were inspected and corrected manually. Reorientation frequency was computed as the inverse of the average time between reorientation events. Reorientation events were detected as described previously [81, 30], by identifying the time points at which both 1) the absolute change in angle exceeded 25° and 2) the filtered velocity was lower than 70% of the average velocity of the trajectory. Videos recorded at 40X were processed with a Fast Radial Symmetry Transform algorithm to remove diffraction rings [82] before applying the tracking routine.

### Phylogenetic tree construction

Phylogenetic tree was constructed using Phylophlan 3.0 [83]. Phylophlan was run according to instructions using all reference genomes for all 4788 strains as the initial dataset. Reference genomes were downloaded from RefSeq. Three outgroups were also added from RefSeq (GCA 000012345, GCA 000168995 and GCA 002355955). Additional references were added using ‘phylophlan get reference’ with the ‘-g c Gammaproteobacteria -n -1’ options. ‘phylophlan write config file’ was run with the ‘-d a – db aa diamond –map dna diamond –map aa diamond –msa mafft –trim trimal –tree1 iqtree’ options and the final phylophlan run was executed using ‘phylophlan’ with the options ‘-d phylophlan –diversity medium –accurate -t a’. The resulting IQTree file was used for phylogenetic analysis and as the basis for the tree in Figure S1, and SH values were added by re-running the dataset in IQTree using the original seed (857918) and resampling 10,000 times.

### Electron microscopy

Flagellation was measured using scanning electron microscopy (SEM). SEM was rendered using an extreme high resolution (XHS) TFS Magellan 400 (ScopeM, ETH Zurich) outfitted with a field emission gun and operated at 2.00 kV and 50 pA. Images of fixed bacteria were obtained using a secondary electron through-the-lens detector. Liquid culture samples were collected at different time points and fixed with 1% (w/v) glutaraldehyde. Samples were then deposited on hydrophilized silicon wafers treated with 0.01% poly-L-lysine. The wafers were successively submerged in 2.5% glutaraldehyde (salinity 27 psu), seawater, 1% osmium tetroxide, and seawater for five min each. This was followed by an ethanol drying series (30%, 50%, 70%, 90%, 100%, with samples submerged for two min for each step), followed by final washing three times in water-free ethanol. The samples were then critical-point dried using the cell-monolayer program (CPD 931 Tousimis, ScopeM), and mounted with silver paint to aluminum stubs. The stubs were then sputter-coated with 4 nm of Pt-Pd (CCU-010 Metal Sputter Coater Safematic, ScopeM) to prevent sample charging. Cell and flagellar length were determined using ImageJ.

### Cell counting and viability measurements

Cell counting was performed by diluting cells by a factor of 100 and staining them with Syber Green (Sigma Aldrich). For samples where the dead fraction was determined, a second sample was stained with with Sytox Green (Thermofischer). Cells were stained at a final concentration of 5 µM for both stains and incubated in the dark for 10 min at room temperature. After staining, cells were counted using a flow cytometer (Beckman Coulter, CytoFLEX S) equipped with a 488 nm laser. Plate counts to determine the CFUs were performed on Marine Broth (1.5% agar) plates with 15 mL liquid per plate. Only plates with 20–350 colonies were included in the analysis.

### Staining of storage granules

DAPI straining to probe polyphosphate levels was based on methods described previously [84, 85]. A 5 mg/mL DAPI (4’,6-diamidino-2-phenylindole dihydrochloride, Thermo Scientific) solution in filtered (milliQ) water, stored at -20 ^*°*^C, was thawed and diluted to 25 µg/mL in ASW as a working stock on the day of the experiment. Samples of 1 mL cell suspension with approximately 1·10^7^cells (OD 0.01) were fixed with 3.7% paraformaldehyde for 1 h and then added to a filter tower pulled through a filter column by pressure difference. The filters (25 mm diameter) consisted of a nitrocellulose backing filter (0.4 µm, Thermo Scientific) covered by a black Isopore (TM) membrane filter (0.2 µm). After filtration, the black membrane filter was placed on 500 µL DAPI solution for 10 min in the dark. The filter was then gently washed by sweeping it through a drop of milliQ water and dried for 10 min. The filter was then placed on a standard glass slide under a 24× 50 mm coverslip, with a small drop (20 µL) of a photostability mixture consisting of 4 parts Citifluor (Citifluor, Ltd.) and 1 part Vectashield (Vector Laboratories, Inc.). Samples were measured using an oil immersion objective (100X, 1.4 NA, Nikon) and a Canon EOS 80D DSLR camera (ISO 800, 0.25 s exposure), with excitation by a broad-spectrum mercury lamp (Prior Scientific) with DAPI filter cube (Chroma, ex: 350/50, di: 400, em: LP420).

The neutral lipid stain Bodipy 493/503 (Thermo Scientific) was used to visualise PHB granules [86]. A stock solution of 1 mg/mL in DMSO was diluted ten fold in DMSO to obtain a working stock of 100 µg/mL. To stain cells, 5 µL dye solution was added to 0.5 mL cell suspension with approximately 1·10^8^ cells/mL (OD 0.1), briefly vortexed and incubated on ice for 10 min in the dark. Cells were then immobilized by placing them on poly-L-lysine (Sigma) coated coverslips for 30 min. Cells were imaged using an oil immersion objective (100X, 1.4 NA, Nikon) and CMOS camera (ORCA Flash 4.0, Hamamatsu) under epifluorescent illumination provided by a mercury lamp (Prior Scientific) with Chroma EGFP filter cube (ex: 470/40 nm, di: 495 nm, em: 525/50 nm). To determine the PHB content per cell, the raw fluorescence signal *F*^*∗*^of a rectangle with one cell was integrated, and the background fluorescence value of an area without cells in the same image subtracted. For each cell, *F*^*∗*^ was then normalized to *F* according to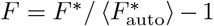, with the autofluorescence 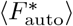value averaged over all cells of a strain without PHB synthesis genes.

### Training and usage of a Naive Bayesian Classifier

Protein-coding sequences from all strains were re-annotated using EGGNOG-mapper [87]. KEGG orthologous group (KOG) assignments [88] from EGGNOG annotations were tabulated for all strains. KOGs were filtered to remove KOGs with representation only from a single strain or KOGs with uniform representation. In the limokinetic classifier, genes with higher relative abundance in the limostatic strains were excluded as potential features (and visa-versa). The feature matrix was then binarized, reducing counts of each KOG to presence/absence data for each strain. Recursive feature elimination was was implemented using the ‘FeatureTerminatoR’ [89] package and attempting training from 2 to 1000 features using ‘leave-one-out’ cross-validation for each strain, assuming a Poisson distribution and using a Laplacian smoothing value of 1. A value of 22 features was chosen as the smallest, high-accuracy feature set that would not be prone to over-fitting (Fig. S11). Training of the final Bayesian classifier was performed by the ‘naivebayes’ [90] and ‘caret’ [91] packages for R. Training was performed using 128 train/test splits, training on 2/3 of the data and reserving 1/3 of the data set for prediction. To test for phylogenetic biases in the classifier outcome, a whole genome phylogeny (Fig. S1) was used in a logistic regression analysis, implemented in the R package phylolm [92].

For the depth profiling using the classifiers, the limokinetic and limostatic classifiers were applied to field data in the Ocean Microbiome Database (OMD1; https://microbiomics.io/ocean, [49]) to profile patterns of occurrence for each phenotype as a function of depth. Feature presence or absence was extracted from pre-calculated KEGG orthlogous groups for each metagenomic assembly (MAG) assembly in the dataset. Abundance was calculated as the coverage of unambiguously (limokinetic positive AND limostatic negative or visa-versa) gammaproteobacterial MAGs divided by the total coverage or total coverage of gammaproteobacterial MAGs. For the normalized depth profiles, the fractional abundance in each sample was log-normalized and mean-centered and scaled independently for each group.

## Acknowledgements

We are grateful to Noele Norris, Gertjan Meijer, Elise Ledieu, Yutaka Yawata, Natasha Blitvic, Vicente Fernandez, and Jing Yan for advice and discussions. We thank Russell Naisbit for scientific editing and Jean-Baptiste Raina for a critical reading of the manuscript. We thank Lucas Paoli and Astrid Stubbush for help with the Ocean Microbiomics Database. We acknowledge Lisa Moor and Ladina Schocher for help with experiments. We thank Knut Drescher, Glen D’Souza, Kirsten Jung, Sophie Brameyer, Matthias Ullrich, Samuel Charlton and Isobel Short for sharing strains. We thank the microscopy facility ScopeM at ETHZ for SEM training, imaging and facilities. This work was supported by the Simons Collaboration on Principles of Microbial Ecosystems (PriME; grants 542395FY22 to R.S. and 542379FY22 to M.A.), the Swiss National Science Foundation (grant 205321 207488 to R.S.) and a Gordon and Betty Moore Foundation Symbiosis in Aquatic Systems Initiative Investigator Award (grant GBMF9197 to R.S.; https://doi.org/10.37807/GBMF9197).

## Author contributions

J.K., F.C., and R.S. conceived the work. J.K., S.Z., and E.C. performed experiments, with help from C.M. J.K. and S.Z. analysed experimental data. Z.L., B.R., and C.M. performed genetic analyses. Z.L. performed model training and testing. All authors discussed and interpreted results. J.K., Z.L., B.R., and R.S. wrote the paper with input from all authors.

## References

[1] F. M. Lauro, D. McDougald, T. Thomas, T. J. Williams, S. Egan, S. Rice, M. Z. DeMaere, L. Ting, H. Ertan, J. Johnson, S. Ferriera, A. Lapidus, I. Anderson, N. Kyrpides, A. C. Munk, C. Detter, C. S. Han, M. V. Brown, F. T. Robb, S. Kjelleberg, and R. Cavicchioli, “The genomic basis of trophic strategy in marine bacteria,” Proceedings of the National Academy of Sciences, vol. 106, pp. 15527–15533, Sept. 2009.

[2] J. P. Zehr, J. S. Weitz, and I. Joint, “How microbes survive in the open ocean,” Science, vol. 357, pp. 646–647, Aug. 2017.

[3] M. Sebastián, M. Estrany, C. Ruiz-González, I. Forn, M. M. Sala, J. M. Gasol, and C. Marrasé, “High Growth Potential of Long-Term Starved Deep Ocean Opportunistic Heterotrophic Bacteria,” Frontiers in Microbiology, vol. 10, p. 760, Apr. 2019.

[4] R. Stocker, “Marine Microbes See a Sea of Gradients,” Science, vol. 338, pp. 628–633, Nov. 2012.

[5] M. Bergkessel, D. W. Basta, and D. K. Newman, “The physiology of growth arrest: uniting molecular and environmental microbiology,” Nature Reviews Microbiology, vol. 14, pp. 549–562, Sept. 2016.

[6] J. Dworkin and C. S. Harwood, “Metabolic Reprogramming and Longevity in Quiescence,” Annual Review of Microbiology, vol. 76, pp. 91–111, Sept. 2022.

[7] B. S. Lambert, V. I. Fernandez, and R. Stocker, “Motility drives bacterial encounter with particles responsible for carbon export throughout the ocean,” Limnology and Oceanography Letters, vol. 4, pp. 113–118, Oct. 2019.

[8] N. Wadhwa and H. C. Berg, “Bacterial motility: machinery and mechanisms,” Nature Reviews Microbiology, Sept. 2021.

[9] J. M. Keegstra, F. Carrara, and R. Stocker, “The ecological roles of bacterial chemotaxis,” Nature Reviews Microbiology, Mar. 2022.

[10] N. I. Wisnoski and J. T. Lennon, “Scaling up and down: movement ecology for microorganisms,” Trends in Microbiology, vol. 31, pp. 242–253, Mar. 2023.

[11] T. M. Hoehler and B. B. Jørgensen, “Microbial life under extreme energy limitation,” Nature Reviews Microbiology, vol. 11, pp. 83–94, Feb. 2013.

[12] C. P. Kempes, P. M. van Bodegom, D. Wolpert, E. Libby, J. Amend, and T. Hoehler, “Drivers of Bacterial Maintenance and Minimal Energy Requirements,” Frontiers in Microbiology, vol. 8, Jan. 2017.

[13] E. Biselli, S. J. Schink, and U. Gerland, “Slower growth of Escherichia coli leads to longer survival in carbon starvation due to a decrease in the maintenance rate,” Molecular Systems Biology, vol. 16, June 2020.

[14] B. Ni, R. Colin, H. Link, R. G. Endres, and V. Sourjik, “Growth-rate dependent resource investment in bacterial motile behavior quantitatively follows potential benefit of chemotaxis,” Proceedings of the National Academy of Sciences, vol. 117, pp. 595–601, Jan. 2020.

[15] M. H. Larsen, N. Blackburn, J. L. Larsen, and J. E. Olsen, “Influences of temperature, salinity and starvation on the motility and chemotactic response of Vibrio anguillarum,” Microbiology, vol. 150, pp. 1283–1290, May 2004.

[16] E. Yam and K. Tang, “Effects of starvation on aggregate colonization and motility of marine bacteria,” Aquatic Microbial Ecology, vol. 48, pp. 207–215, Aug. 2007.

[17] M. A. Lever, K. L. Rogers, K. G. Lloyd, J. Overmann, B. Schink, R. K. Thauer, T. M. Hoehler, and B. B. Jørgensen, “Life under extreme energy limitation: a synthesis of laboratory- and field-based investigations,” FEMS Microbiology Reviews, vol. 39, pp. 688–728, Sept. 2015.

[18] X. Wei and W. D. Bauer, “Starvation-Induced Changes in Motility, Chemotaxis, and Flagellation of Rhizobium meliloti,” Applied and Environmental Microbiology, vol. 64, pp. 1708–1714, May 1998.

[19] K. Malmcrona-Friberg, A. Goodman, and S. Kjelleberg, “Chemotactic Responses of Marine Vibrio sp. Strain S14 (CCUG 15956) to Low-Molecular-Weight Substances under Starvation and Recovery Conditions,” Applied and Environmental Microbiology, vol. 56, pp. 3699–3704, Dec. 1990.

[20] X. Zhuang, S. Guo, Z. Li, Z. Zhao, S. Kojima, M. Homma, P. Wang, C. Lo, and F. Bai, “Live-cell fluorescence imaging reveals dynamic production and loss of bacterial flagella,” Molecular Microbiology, vol. 114, pp. 279–291, Aug. 2020.

[21] J. L. Ferreira, F. Z. Gao, F. M. Rossmann, A. Nans, S. Brenzinger, R. Hosseini, A. Wilson, A. Briegel, K. M. Thormann, P. B. Rosenthal, and M. Beeby, “Gammaproteobacteria eject their polar flagella under nutrient depletion, retaining flagellar motor relic structures,” PLOS Biology, vol. 17, p. e3000165, Mar. 2019.

[22] E. Ledieu-Dhebércourt, Lost in Starvation: How the interplay between physiology and ecology impacts bacterial persistence in a patchy landscape. PhD thesis, MIT, Cambridge, USA, Sept. 2022.

[23] S. Stretton, S. J. Danon, S. Kjelleberg, and A. E. Goodman, “Changes in cell morphology and motility in the marine Vibrio sp. strain S14 during conditions of starvation and recovery,” FEMS Microbiology Letters, vol. 146, pp. 23–29, Jan. 2006.

[24] J.-B. Raina, B. S. Lambert, D. H. Parks, C. Rinke, N. Siboni, A. Bramucci, M. Ostrowski, B. Signal, A. Lutz, H. Mendis, F. Rubino, V. I. Fernandez, R. Stocker, P. Hugenholtz, G. W. Tyson, and J. R. Seymour, “Chemotaxis shapes the microscale organization of the ocean’s microbiome,” Nature, vol. 605, pp. 132–138, May 2022.

[25] U. Alcolombri, F. J. Peaudecerf, V. I. Fernandez, L. Behrendt, K. S. Lee, and R. Stocker, “Sinking enhances the degradation of organic particles by marine bacteria,” Nature Geoscience, vol. 14, pp. 775–780, Sept. 2021.

[26] T. Kiørboe, A mechanistic approach to plankton ecology. Princeton University Press, 2008.

[27] J. Slomka, U. Alcolombri, E. Secchi, R. Stocker, and V. I. Fernandez, “Encounter rates between bacteria and small sinking particles,” New Journal of Physics, vol. 22, p. 043016, Apr. 2020.

[28] B. Borer, I. H. Zhang, A. E. Baker, G. A. O’Toole, and A. R. Babbin, “Porous marine snow differentially benefits chemotactic, motile, and nonmotile bacteria,” PNAS Nexus, vol. 2, p. pgac311, Feb. 2023.

[29] T. M. Steinum, S. Karataş, N. T. Martinussen, P. M. Meirelles, F. L. Thompson, and D. J. Colquhoun, “Multilocus Sequence Analysis of Close Relatives Vibrio anguillarum and Vibrio ordalii,” Applied and Environmental Microbiology, vol. 82, pp. 5496–5504, Sept. 2016.

[30] K. Son, J. S. Guasto, and R. Stocker, “Bacteria can exploit a flagellar buckling instability to change direction,” Nature Physics, vol. 9, pp. 494–498, Aug. 2013.

[31] L. Xie, T. Altindal, S. Chattopadhyay, and X.-L. Wu, “Bacterial flagellum as a propeller and as a rudder for efficient chemotaxis,” Proceedings of the National Academy of Sciences, vol. 108, pp. 2246–2251, Feb. 2011.

[32] E. Leifson, B. J. Cosenza, R. Miuirichelano, and I. C. Cleverdon, “Motile marine bacteria. Techniques, ecology and general characteristics,” Journal of Bacteriology, vol. 87, pp. 652–666, 1964.

[33] T. T. Renault, A. O. Abraham, T. Bergmiller, G. Paradis, S. Rainville, E. Charpentier, C. C. Guet, Y. Tu, K. Namba, J. P. Keener, T. Minamino, and M. Erhardt, “Bacterial flagella grow through an injection-diffusion mechanism,” eLife, vol. 6, p. e23136, Mar. 2017.

[34] L. Turner, A. S. Stern, and H. C. Berg, “Growth of Flagellar Filaments of Escherichia coli Is Independent of Filament Length,” Journal of Bacteriology, vol. 194, pp. 2437–2442, May 2012.

[35] X.-Y. Zhuang and C.-J. Lo, “Construction and Loss of Bacterial Flagellar Filaments,” Biomolecules, vol. 10, p. 1528, Nov. 2020.

[36] M. Chen, Z. Zhao, J. Yang, K. Peng, M. A. Baker, F. Bai, and C.-J. Lo, “Length-dependent flagellar growth of Vibrio alginolyticus revealed by real time fluorescent imaging,” eLife, vol. 6, p. e22140, Jan. 2017.

[37] S. Chattopadhyay, R. Moldovan, C. Yeung, and X. L. Wu, “Swimming efficiency of bacterium Escherichia coli,” Proceedings of the National Academy of Sciences, vol. 103, pp. 13712–13717, Sept. 2006.

[38] J. R. Taylor and R. Stocker, “Trade-Offs of Chemotactic Foraging in Turbulent Water,” Science, vol. 338, pp. 675–679, Nov. 2012.

[39] J. G. Mitchell, “The influence of cell size on marine bacterial motility and energetics,” Microbial Ecology, vol. 22, pp. 227–238, Dec. 1991.

[40] N. Cermak, J. W. Becker, S. M. Knudsen, S. W. Chisholm, S. R. Manalis, and M. F. Polz, “Direct single-cell biomass estimates for marine bacteria via Archimedes’ principle,” The ISME Journal, vol. 11, pp. 825–828, Mar. 2017.

[41] B. R. K. Roller, C. Hellerschmied, Y. Wu, T. P. Miettinen, A. L. Gomez, S. R. Manalis, and M. F. Polz, “Single-cell mass distributions reveal simple rules for achieving steady-state growth,” mBio, vol. 14, pp. e01585–23, Oct. 2023.

[42] W. R. Shoemaker, S. E. Jones, M. E. Muscarella, M. G. Behringer, B. K. Lehmkuhl, and J. T. Lennon, “Microbial population dynamics and evolutionary outcomes under extreme energy limitation,” Proceedings of the National Academy of Sciences, vol. 118, p. e2101691118, Aug. 2021.

[43] S. J. Schink, E. Biselli, C. Ammar, and U. Gerland, “Death Rate of E. coli during Starvation Is Set by Maintenance Cost and Biomass Recycling,” Cell Systems, vol. 9, pp. 64–73.e3, July 2019.

[44] D. Kadouri, E. Jurkevitch, and Y. Okon, “Involvement of the Reserve Material Poly-Hydroxybutyrate in Azospirillum brasilense Stress Endurance and Root Colonization,” Applied and Environmental Microbiology, vol. 69, p. 7, 2003.

[45] M. H. Rashid, N. N. Rao, and A. Kornberg, “Inorganic Polyphosphate Is Required for Motility of Bacterial Pathogens,” Journal of Bacteriology, vol. 182, pp. 225–227, Jan. 2000.

[46] K. Sekar, S. M. Linker, J. Nguyen, A. Grünhagen, R. Stocker, and U. Sauer, “Bacterial Glycogen Provides Short-Term Benefits in Changing Environments,” Applied and Environmental Microbiology, vol. 86, pp. e00049–20, Apr. 2020.

[47] L. Bourassa and A. Camilli, “Glycogen contributes to the environmental persistence and transmission of Vibrio cholerae,” Molecular Microbiology, vol. 72, pp. 124–138, Mar. 2009.

[48] J. J. Morris, A. L. Rose, and Z. Lu, “Reactive oxygen species in the world ocean and their impacts on marine ecosystems,” Redox Biology, vol. 52, p. 102285, June 2022.

[49] L. Paoli, H.-J. Ruscheweyh, C. C. Forneris, F. Hubrich, S. Kautsar, A. Bhushan, A. Lotti, Q. Clayssen, G. Salazar, A. Milanese, C. I. Carlström, C. Papadopoulou, D. Gehrig, M. Karasikov, H. Mustafa, M. Larralde, L. M. Carroll, P. Sánchez, A. A. Zayed, D. R. Cronin, S. G. Acinas, P. Bork, C. Bowler, T. O. Delmont, J. M. Gasol, A. D. Gossert, A. Kahles, M. B. Sullivan, P. Wincker, G. Zeller, S. L. Robinson, J. Piel, and S. Sunagawa, “Biosynthetic potential of the global ocean microbiome,” Nature, vol. 607, pp. 111–118, July 2022.

[50] C. D. Amsler, M. Cho, and P. Matsumura, “Multiple factors underlying the maximum motility of Escherichia coli as cultures enter post-exponential growth.,” Journal of Bacteriology, vol. 175, pp. 6238–6244, Oct. 1993.

[51] J. G. Mitchell, L. Pearson, A. Bonazinga, S. Dillon, H. Khouri, and R. Paxinos, “Long lag times and high velocities in the motility of natural assemblages of marine bacteria.,” Applied and environmental microbiology, vol. 61, no. 3, pp. 877–882, 1995.

[52] T. Honda, J. Cremer, L. Mancini, Z. Zhang, T. Pilizota, and T. Hwa, “Coordination of gene expression with cell size enables Escherichia coli to efficiently maintain motility across conditions,” Proceedings of the National Academy of Sciences, vol. 119, p. e2110342119, Sept. 2022.

[53] J. Cremer, T. Honda, Y. Tang, J. Wong-Ng, M. Vergassola, and T. Hwa, “Chemotaxis as a navigation strategy to boost range expansion,” Nature, vol. 575, pp. 658–663, Nov. 2019.

[54] S. Gude, E. Pinçe, K. M. Taute, A.-B. Seinen, T. S. Shimizu, and S. J. Tans, “Bacterial coexistence driven by motility and spatial competition,” Nature, vol. 578, pp. 588–592, Feb. 2020.

[55] P. S. Amy and R. Y. Morita, “Starvation-Survival Patterns of Sixteen Freshly Isolated Open-Ocean Bacteria,” Applied and Environmental Microbiology, vol. 45, pp. 1109–1115, Mar. 1983.

[56] J. Taktikos, H. Stark, and V. Zaburdaev, “How the Motility Pattern of Bacteria Affects Their Dispersal and Chemotaxis,” PLoS ONE, vol. 8, p. e81936, Dec. 2013.

[57] N. W. Frankel, W. Pontius, Y. S. Dufour, J. Long, L. Hernandez-Nunez, and T. Emonet, “Adaptability of non-genetic diversity in bacterial chemotaxis,” eLife, vol. 3, Oct. 2014.

[58] D. R. Brumley, F. Carrara, A. M. Hein, Y. Yawata, S. A. Levin, and R. Stocker, “Bacteria push the limits of chemotactic precision to navigate dynamic chemical gradients,” Proceedings of the National Academy of Sciences, vol. 116, pp. 10792–10797, May 2019.

[59] K. Son, F. Menolascina, and R. Stocker, “Speed-dependent chemotactic precision in marine bacteria,” Proceedings of the National Academy of Sciences, vol. 113, pp. 8624–8629, Aug. 2016.

[60] Y. Yawata, F. Carrara, F. Menolascina, and R. Stocker, “Constrained optimal foraging by marine bacterioplankton on particulate organic matter,” Proceedings of the National Academy of Sciences, vol. 117, pp. 25571–25579, Oct. 2020.

[61] M. Grognot, A. Mittal, M. Mah’moud, and K. M. Taute, “Vibrio cholerae Motility in Aquatic and Mucus-Mimicking Environments,” Applied and Environmental Microbiology, vol. 87, pp. e01293–21, Sept. 2021.

[62] R. Sathyamoorthy, Y. Kushmaro, O. Rotem, O. Matan, D. E. Kadouri, A. Huppert, and E. Jurkevitch, “To hunt or to rest: prey depletion induces a novel starvation survival strategy in bacterial predators,” The ISME Journal, vol. 15, pp. 109–123, Sept. 2020.

[63] S. Subramanian and D. B. Kearns, “Functional Regulators of Bacterial Flagella,” Annual Review of Microbiology, vol. 73, pp. 225–246, Sept. 2019.

[64] S. Zhu and B. Gao, “Bacterial Flagella Loss under Starvation,” Trends in Microbiology, vol. 28, pp. 785–788, Oct. 2020.

[65] J. Pernthaler, “Predation on prokaryotes in the water column and its ecological implications,” Nature Reviews Microbiology, vol. 3, pp. 537–546, July 2005.

[66] C. Matz and K. Jürgens, “High Motility Reduces Grazing Mortality of Planktonic Bacteria,” Applied and Environmental Microbiology, vol. 71, pp. 921–929, Feb. 2005.

[67] F. de Schaetzen, M. Fan, U. Alcolombri, F. J. Peaudecerf, D. Drissner, M. J. Loessner, R. Stocker, and M. Schuppler, “Random encounters and amoeba locomotion drive the predation of Listeria monocytogenes by Acanthamoeba castellanii,” Proceedings of the National Academy of Sciences, vol. 119, p. e2122659119, Aug. 2022.

[68] R. Lewin, “Saprospira grandis: A Flexibacterium That Can Catch Bacterial Prey by “Ixotrophy”,” Microbial Ecology, vol. 34, pp. 232–236, Nov. 1997.

[69] S. Z. Schade, J. Adler, and H. Ris, “How Bacteriophage x Attacks Motile Bacteria,” Journal of Virology, vol. 1, no. 3, pp. 599–609, 1967.

[70] J. Y. Yen, K. M. Broadway, and B. E. Scharf, “Minimum Requirements of Flagellation and Motility for Infection of Agrobacterium sp. Strain H13-3 by Flagellotropic Bacteriophage 7-7-1,” Applied and Environmental Microbiology, vol. 78, pp. 7216–7222, Oct. 2012.

[71] D. E. Hunt, L. A. David, D. Gevers, S. P. Preheim, E. J. Alm, and M. F. Polz, “Resource Partitioning and Sympatric Differentiation Among Closely Related Bacterioplankton,” Science, vol. 320, pp. 1081–1085, May 2008.

[72] O. X. Cordero, H. Wildschutte, B. Kirkup, S. Proehl, L. Ngo, F. Hussain, F. Le Roux, T. Mincer, and M. F. Polz, “Ecological Populations of Bacteria Act as Socially Cohesive Units of Antibiotic Production and Resistance,” Science, vol. 337, pp. 1228–1231, Sept. 2012.

[73] Y. Yawata, O. X. Cordero, F. Menolascina, J.-H. Hehemann, M. F. Polz, and R. Stocker, “Competition-dispersal tradeoff ecologically differentiates recently speciated marine bacterioplankton populations,” Proceedings of the National Academy of Sciences, vol. 111, pp. 5622–5627, Apr. 2014.

[74] J.-H. Hehemann, P. Arevalo, M. S. Datta, X. Yu, C. H. Corzett, A. Henschel, S. P. Preheim, S. Timberlake, E. J. Alm, and M. F. Polz, “Adaptive radiation by waves of gene transfer leads to fine-scale resource partitioning in marine microbes,” Nature Communications, vol. 7, p. 12860, Sept. 2016.

[75] M. S. Datta, E. Sliwerska, J. Gore, M. F. Polz, and O. X. Cordero, “Microbial interactions lead to rapid micro-scale successions on model marine particles,” Nature Communications, vol. 7, p. 11965, June 2016.

[76] B.-H. Y. and R. E., “A novel Vibrio sp. pathogen of the coral Pocillopora damicornis,” Marine Biology, vol. 141, pp. 47–55, July 2002.

[77] R. M. Welsh, J. R. Zaneveld, S. M. Rosales, J. P. Payet, D. E. Burkepile, and R. V. Thurber, “Bacterial predation in a marine host-associated microbiome,” The ISME Journal, vol. 10, pp. 1540–1544, June 2016.

[78] R. Stocker, J. R. Seymour, A. Samadani, D. E. Hunt, and M. F. Polz, “Rapid chemotactic response enables marine bacteria to exploit ephemeral microscale nutrient patches,” Proceedings of the National Academy of Sciences, vol. 105, pp. 4209–4214, Mar. 2008.

[79] E. C. Kaeppel, A. Gärdes, S. Seebah, H.-P. Grossart, and M. S. Ullrich, “Marinobacter adhaerens sp. nov., isolated from marine aggregates formed with the diatom Thalassiosira weissflogii,” International Journal of Systematic and Evolutionary Microbiology, vol. 62, pp. 124–128, Jan. 2012.

[80] D. B. Allen, T. Caswell, N. C. Keim, C. M. van der Wel, and R. W. Verweij, “Trackpy,” 2021. 10.5281/zenodo.4682814.

[81] E. E. Clerc and et al., “Strong chemotaxis by marine bacteria towards polysaccharides is enhanced by the abundant organosulfur compound DMSP,” Nature Communications, vol. 14, p. 8080, 2023.

[82] G. Loy and A. Zelinsky, “A Fast Radial Symmetry Transform for Detecting Points of Interest,” Computer Vision ECCV 2002, vol. 2350, pp. 358–368, 2002. Series Title: Lecture Notes in Computer Science.

[83] F. Asnicar, A. M. Thomas, F. Beghini, C. Mengoni, S. Manara, P. Manghi, Q. Zhu, M. Bolzan, F. Cumbo, U. May, J. G. Sanders, M. Zolfo, E. Kopylova, E. Pasolli, R. Knight, S. Mirarab, C. Huttenhower, and N. Segata, “Precise phylogenetic analysis of microbial isolates and genomes from metagenomes using PhyloPhlAn 3.0,” Nature Communications, vol. 11, p. 2500, May 2020.

[84] M. Streichan, J. R. Golecki, and G. SchÃ¶n, “Polyphosphate-accumulating bacteria from sewage plants with different proceses for biological phosphorus removal,” FEMS Microbiology Letters, vol. 73, pp. 113–124, Feb. 1990.

[85] D. P. Mesquita, A. L. Amaral, C. Leal, M. Carvalheira, J. R. Cunha, A. Oehmen, M. A. M. Reis, and E. C. Ferreira, “Monitoring intracellular polyphosphate accumulation in enhanced biological phosphorus removal systems by quantitative image analysis,” Water Science and Technology, vol. 69, pp. 2315–2323, June 2014.

[86] J. Kacmar, R. Carlson, S. J. Balogh, and F. Srienc, “Staining and quantification of poly-3-hydroxybutyrate in Saccharomyces cerevisiae and Cupriavidus necator B cell populations using automated flow cytometry,” Cytometry Part A, vol. 69A, pp. 27–35, Jan. 2006.

[87] A. Hernández-Plaza, D. Szklarczyk, J. Botas, C. Cantalapiedra, J. Giner-Lamia, D. R. Mende, R. Kirsch, T. Rattei, I. Letunic, L. Jensen, P. Bork, C. von Mering, and J. Huerta-Cepas, “eggNOG 6.0: enabling comparative genomics across 12 535 organisms,” Nucleic Acids Research, vol. 51, pp. D389–D394, Jan. 2023.

[88] M. Kanehisa and S. Goto, “KEGG: Kyoto Encyclopedia of Genes and Genomes,” Nucleic Acids Research, vol. 28, No. 1, pp. 27–30, 2000.

[89] G. Hutson, “Feature Selection Engine to Remove Features with Minimal Predictive Power,” 2022.

[90] M. Majka, “naivebayes: High Performance Implementation of the Naive Bayes Algorithm in R,” 2019.

[91] M. Kuhn, “Building Predictive Models in R Using the caret Package,” Journal of Statistical Software, vol. 28, pp. 1–26, Nov. 2008. 10.18637/jss.v028.i05.

[92] L. S. Tung Ho and C. Ané, “A Linear-Time Algorithm for Gaussian and Non-Gaussian Trait Evolution Models,” Systematic Biology, vol. 63, pp. 397–408, May 2014.

