## Supplementary Information for "Risk-reward trade-off in motility endurance generates dichotomy in search strategies among copiotrophic marine bacteria"

February 26, 2025

**3 Supplementary Figures** **8**

**4 Supplementary Tables** **22**

**5 Supplementaty Videos** **26**

### 1 Supplementary Text

#### 1.1 Excluding the effect of alternative energy sources for motility

##### 1.1.1 Necromass recycling

We considered the recycling of necromass, the biomass of dead cells, as a potential mechanism providing energy for motility. Necromass recycling has been shown to be a possible mechanism to increase the survival of bacterial populations under prolonged starvation by attenuating the death rate of cells [56, 53]. However, in our starvation experiments the number of cells does not decrease over time for any of the strains, indicating that the net cell death, if any, is small (Fig. 3C). We cannot rule out that there is some initial cell death that is masked by the increase in the number of cells due to reductive divisions. Therefore, we measured cell death to estimate how much necromass is released during starvation.

Direct measurements of cell death using live/dead staining in the limokinetic and limostatic strains revealed overall a small difference in the fraction of dead cells between the two classes (Fig. S8E). The fraction of dead cells decreased over time, which shows that necromass does not accumulate over the course of the starvation time and indicating that cell lysis occurs primarily at the onset of starvation. At the onset of starvation the average fraction of dead cells was $0.12 \pm 0.04$ and $0.23 \pm 0.03$ (mean $\pm$ one standard error of the mean), respectively, for limostatic and limokinetic strains. The difference between limostatic and limokinetic strains suggests limokinetic strains release 11 % more biomass in the form of necromass, which could in principle be available for swimming and fuel motility for approximately one day. However, this amount of biomass is much lower than decrease in biomass observed in limokinetic strains of 62 % over one week, showing that necromass does not act as long-term energy source. Furthermore, the efficiency of biomass recycling ($\sim$20%, estimated in starvation experiments using *E. coli* [53]), indicates that the actual amount of necromass that can be re-absorbed into the population is even lower. We conclude that in our experiments biomass recycling plays at most a minor role in supplying cells with energy for motility.

##### 1.1.2 Light

A potential alternative energy source during carbon starvation is light, which some bacteria can capture using proteorhodopsins. This strategy is widely used by oligotrophic non-motile bacteria, like SAR11 [21, 59]. Proteorhodopsins have also been proposed to provide energy for motility, a process that has been demonstrated in engineered (proton-motorized) *E. coli* [65]. Light-driven sodium pumps [28], a special class of rhodopsins, could potentially enable (sodium-motorized) marine bacteria to fuel motility using light. However, none of the 26 strains tested possesses proteorhodopsin genes. In addition, motility levels of *V. anguillarum* FS144 and *V. coralliilyticus* YB1 in the dark did not differ from those in the light (Fig. S9A). Together, these data indicate that the energy source for limokinetic cells during starvation does not come from light.

##### 1.1.3 Trace levels of nutrients in starvation medium

Additional experiments confirmed that the energy source that powers motility under starvation is internal to the cells. The chemical energy for swimming could be external to the cells (e.g., provided by low levels of residual nutrients in the medium) or internal (e.g., resulting from the conversion of biomass to energy). These two sources would be expected to yield different relationships between the motility level and the concentration of cells. An external energy source implies decreasing energy for motility per cell with increasing cell concentration, as cells must share the available nutrients, whereas for an internal energy source the energy for motility per cell would be independent of the cell concentration. To distinguish between these two possibilities, we measured the dependence of motility on cell concentration for four limokinetic strains under starvation. We prepared a dilution series of cell concentration spanning two orders of magnitude and quantified the fraction of motile cells for at least 24 h following the onset of starvation. A linear fit to the motile fraction as a function of the logarithm of the relative cell concentration reveals a positive slope for all strains, with average slope of $0.08 \pm 0.04$ (Fig. S9B), thereby excluding a negative dependence of the motile fraction on cell concentration. Together with the decrease in biomass (Fig 3), this result shows that the energy source to power swimming during starvation in limokinetic cells is internal to the cells.

#### 1.2 Function of the selected features for genomic classification

##### 1.2.1 Resource conservation and fatty acid metabolism

A small number of genes associated with limokinetic strains are involved in energy or resource conservation. Limokinetic strains convert biomass to energy to fuel motility (Main text, Fig. 3). A possible energy source for cells are lipids, a high-energy component of cellular biomass. A number of accessory genes involved with fatty acid degradation were selected for the classifier. The presence of malonate semialdehyde dehydrogenase (K00140; *mmsA*) and inorganic pyrophosphatase (K01507; *ppa*) seems to indicate an important role for efficient fatty acid degradation in these cells, as both are downstream metabolites formed from the undesirable leftovers of beta-oxidation (propanoyl-CoA and pyrophosphate) and these enzymes help to confer complete degradation of these byproducts. The presence of a short-chain fatty acid transporter (K02106,*atoE*) and glyoxylate/hydroxypyruvate reductase (K12972; *ghrA*) could also allow for the uptake and use of small carboxylic acids from the water column. The selected set also includes electron transfer flavoproteins (K03521; *etfA*), which efficiently couple the production of $FADH_2$ from beta-oxidation to the ubiquinone pool of the electron transport chain, linking fatty acid degradation directly to the production of ATP [66].

##### 1.2.2 Oxidative stress defense

Many features of our classifier relate to reactive oxygen species (ROS) and related oxidative stress. ROS [26] are abundant in marine environments [3, 43] and lead to oxidative stress, which is believed to be especially detrimental for cells in situations that permit limited or no growth [39, 45], because the ability to regenerate damaged proteins or cellular machinery is limited. Limokinetic strains are likely experiencing greater oxidative stress during starvation than limostatic strains, as motility requires an increased metabolism and respiration level, which in turn generates ROS and increases ROS-associated cell damage [8].

The strongest direct evidence for a strategy of mitigating oxidative stress is the presence of catalase-peroxidase *katG* (K03782) as one of the most highly weighted features our set associated with limokinetic strains. Another protein in our feature set, Dps (K04047), is a DNA-binding protein that is typically expressed under starvation conditions and non-specifically binds DNA, forming a ROS-resistant structure [7, 10, 37]. Other indirect or systemic adaptations are also present, such as phosphogluconate dehydratase (K01690; *edd*), indicating a preference for the Entner–Doudoroff pathway (EDP), for which it acts as the entry point. The EDP has been shown to be preferentially used over the Embden–Meyerhof–Parnas pathway (EMP) in a number of marine bacteria as it appears to confer additional resistance to oxidative stress [33]. K09797 is also present and represents a protein of unknown function, but homologous proteins have been annotated as 'oxidative stress response proteins' in a number of other bacteria.

To test if the two groups of strains showed different sensitivities to ROS stress, we measured the effect of ROS on selected strains from each group by exposing them to hydrogen peroxide. When we applied hydrogen peroxide during exponential growth, we measured no significant difference between limokinetic and limostatic strains in the growth rate (Fig. S12A) or motility level (Fig. S12B). We then applied different levels of hydrogen peroxide oxidative stress to cells starved for 24 h and measured the lag time of growth resumption after nutrients were once again provided (Fig. S12C). These experiments revealed a mildly elevated increase in sensitivity of limostatic cells to oxidative stress compared to the lag time without added hydrogen peroxide (on average 1.36-fold vs. 1.09-fold, Fig S12C), but the oxidative stress level ($[H_2O_2]$ = 10 μM) at which this effect occurred was higher than cells would experience in nature. Typically, oxidative stress in marine environments is expected to be low in intensity ($[H_2O_2] \approx$100 nM) but long-lasting [27]. Therefore, it is possible that differences between the two classes are only apparent when the effects of oxidative stress accumulate on longer timescales than investigated in this study, for example through DNA and protein damage.

Finally, it is possible that the presence of ROS-related genes are a reflection of the community context of the bacterial cells. It has been shown that cyanobacteria can 'offload' the cost of ROS detoxification to algal cells that have to perform this function anyway [41], an example of the black queen hypothesis [42]. As limostatic strains rapidly lose motility, they likely prevail in situations where the distance between particles is small, such as algal blooms, where the community provides protection against ROS. By contrast, limokinetic strains, must provide their own ROS detoxification machinery in order to survive in scenarios with low particle (and algal) concentrations.

##### 1.2.3 Cysteine and methionine synthesis

Another series of genes associated with the limokinetic response are involved in synthesis of sulfur-containing amino acids, such as methionine and cysteine. MetX (K00641) catalyzes the synthesis of the homocysteine and methionine precursor acetyl-L-homoserine. This is less common alternative to the more conventional route via succinyl-L-homoserine commonly found in Gammaproteobacteria [16]. Reasons for preferring one pathway over the other are not known, but it could be speculated that the use of the acetyl-L-homoserine pathway constitutes a slight fitness advantage during starvation, as it requires fewer enzymes and is able to incorporate sulfide directly without the use of a cysteine intermediate (cystathionine). Also associated with homocysteine and methionine metabolism, the *cysG* (K02302) gene encodes a multifunctional enzyme that acts as a methyltransferase implicated in both sirohaem and vitamin B12 synthesis [20]. Importantly, sirohaem is a required cofactor for assimilatory sulfite reductases that produce hydrogen sulfide to be used in homocysteine and methionine synthesis [62]. The *cysG* gene also functions in cobalamin (B12) maturation and its presence may represent the importance of B12 synthesis in this regard [20] - methylation of homocysteine is the final step in methionine biosynthesis and B12-utilizing enzymes (metH) for this reaction have been shown to have a $\sim$30-fold increase in activity over B12-independent proteins[4]. Overall, the presence of *cysG* and related genes is likely due to an overall heightened requirement for sulfur-containing amino acids. One possibility for an increased demand of these amino acids is a demand for methionine to act as a methyl-donating molecule in chemotactic cells [11]. Another possibility is that this is also related to protein repair in response to oxidative stress, as cysteine and methionine are reported to be most sensitive to ROS damage [15]. Cysteine is also a precursor for the production of glutathione, the main ROS-scavenging molecule in the cell, suggesting that the genes involved in cysteine and methionine synthesis could play additional roles in oxidative stress defense.

##### 1.2.4 Motility and chemotaxis

A number of motility-related regulatory genes are implicated as features of the limokinetic strains. Included among these are two-component regulatory systems for the expression of flagella under quorum sensing [58] (K07666, K07645; *qseB, qseC*) or swarming [22] (K20977, K20978; *hsbR, hsbA*) scenarios. It is likely that these are in some way responsible for triggering the decision to swim in these cells and they could be potential targets for future studies into the mechanism underlying the limokinetic–limostatic distinction. Accessory proteins such as CheC and CheD (K03410,K03411) are also present. In *B. subtilis*, CheC and CheD have been shown to form an alternative mechanism of sensory adaptation that works alongside methylation-dependent adaptation (albeit with a likely lower dynamic range) [49]. Sensory adaptation through methylation is an energetically expensive trait, equivalent to approximately 10% of the cost of turning the flagellar motor [35]. This means that cells might favor using CheCD for sensory adaptation in situations of starvation, where energy or methyl-group donor molecules (like methionine, see 'Cysteine and methionine synthesis' above) are scarce [11].

##### 1.2.5 Capsule and biofilm formation

Capsular polysaccharides can protect against adverse environmental conditions including desiccation or osmotic stress, unfavorable pH, the presence of antibacterial compounds, viral predation, and oxidative stress, and are common in marine bacteria [60]. The production of capsular polysaccharides has been shown to confer fitness advantages in gammaproteobacteria under nutrient-limited conditions [5]. The *vpsO, vpsM* and *vpsN* genes involved in capsular polysaccharide synthesis and export are present in our final set of classifier features (K16554, K20920, K20988), as is *wcaJ* (undecaprenyl-phosphate glucose phospotransferase; K03606), which is required for the first step in capsule formation, linking sugar residues to a lipid anchoring moiety [54, 17]. Also related to capsule formation, CDP-4-dehydro-6-deoxyglucose reductase (K00523) produces precursors for O-antigen/Lipopolysaccharide production, which has been shown to confer resistance to a variety of environmental stressors [19, 70].

Several capsule polysaccharide genes are also involved in extracellular polysaccharide production of Vibrio (exo)polysaccharide (VPS) biofilms [68]. Within the Vibrionaceae, the possession of VPS genes can be used as a predictor of limokinetic behavior (Table S4). Interestingly, we discovered that the VPS-assocated gene rbmC (not part of the classifier) can correctly predict limokinetic behavior within the Vibrionaceae (accuracy 93%, when including all the Vibrio species in this study; Table S4). RbmC has been shown to facilitate linking of VPS-biofilm to host-produced glycans such as mucus [23], and could thereby assist host colonization by marine bacteria. This suggests limokinetic behavior could play a role in pathogenicity and symbiosis [38, 48].

#### 1.3 Phylogenomic signature in the classifier

We investigated a potential bias in our classifier due to phylogenetic correlations in the data, and focused first if the feature (Orthologous group) could be explained by phylogeny. This was done to examine if the patterns we observed are due to only closely related bacteria having similar traits, rather than a broader association across the more divergent bacteria in our dataset. First, the binary outcome of both the limokinetic as well as the limostatic classifier were regressed against each individual KEGG ortholog presence/absence in all genomes using a generalized linear model that did not consider phylogeny in the logistic regression. We then included a whole genome phylogeny in a separate phylogenetic generalized linear model framework to assess if phylogeny could explain any observed association as implemented in the R package phylolm [63] by comparing the statistical significance of the slope coefficient in the two regression models.

One well-known disadvantage of logistic regression occurs when near-complete separation of genomic features and the classification outcome occur, leading to a large uncertainty of the coefficient estimates [6], as is the case for, for example, K07666 (Fig. 5A). Therefore, features where the logistic regression without including phylogeny were insignificant were further examined with alternative phylogentic comparative methods: Pagel's method for correlated binary trait evolution [46] as implemented in the phytools R package [50] and phylogenetic linear regression on the binary variables using Pagel's lambda as implemented in the R package phylolm [63]. These alternative methods confirmed K07666 is a robust feature of the dataset, but the limokinetic classifier contains four features where an influence of phylogeny cannot be excluded (K02106,K12972,K03521,K09797) (Table S3).

Finally, we performed a complimentary analysis by combining all of the genomic features using phylogenetic principal components analysis [50] and assessed if the orthogonal combinations of features in the classifier (principal component scores for each genome) were associated with the Bayesian classifier outcome using the same phylogenetic logistic regression approach outlined above. For the limokinetic classifier, the first 6 principal component axes of phylogenetic PCA explain 89.2 % of the total variation in the dataset. PC1 explains 39.3 % of total variation and has the highest level of significance when including phylogeny (p=0.08). The 5 strongest loadings on this principal component in order are K07666, K07645, K01507, K03411, K20977. For the limostatic classifier, the first 6 principal component axes of phylogenetic PCA explain 72.1 % of the total variation in the dataset. PC1 explains 27.8 % of total variation and is highly significant when included for phylogeny (p=0.003) and the 5 strongest loadings (an arbitrary cutoff) on this principal component in order are: K11922, K12113, K11198, K11199, K11200. This further confirms that these classifier features cannot be explained by phylogeny alone.

We then investigated the consequences of phylogenetic bias in classifier prediction. The collection used to train the classifier consists of 19 strains from the Vibrionaceae family (the genera *Vibrio* and *Aliivibrio*), of which the majority (14) are limostatic, and 7 strains from other families (3 Alteromonadaceae, 2 Oceanospirillaceae and 2 Phyllobacteriaceae), of which the majority (6) are limokinetic (Fig. S1, Table S1). Therefore, a taxonomic model that classifies all Vibrionaceae strains as limostatic and all others as limokinetic would accurately predict the behavior of 20/26 strains (77%), significantly different from a binomial random classifier ($p = 0.004$, calculated as the probability of correctly predicting at least 20 of 26 strains, assuming a binomial distribution with 50% as the chance of a successful prediction for each strain). If we include the 6 additional strains used for prediction, this decreases slightly to 24/32 correct (75%). The Bayesian limokinetic and limostatic classifiers built in this work (Fig. 5) both correctly predict 23/26 and 28/32 of the strains (88%), outperforming the phylogenetic classifier ($p = 0.07$, the chance of predicting at least 28 out of 32 strains correctly, with a success chance for each strain of 75%). Within all Vibrionaceae in this study, the classifier correctly predicts 20/23 strains. This is better than a phylogenetic approach ($p = 0.05$, based on the chance of predicting at least 20/23 correctly with an individual chance of 16/23 = 69.5%). Hence, in this dataset there appears to be a taxonomic component associated with the limokinetic/limostatic dichotomy, but this classifies the strains less accurately than a Bayesian classifier built upon genomic information.

### 2 Supplemental discussion

#### 2.1 How cells can optimize their exploratory behavior

The encounter rate of bacteria with larger particles scales linearly with the effective diffusion coefficient that describes the bacteria's random walk [32]. Hence, the effective diffusion coefficient of bacteria is a quantitative measure of how much space they explore. The average diffusion coefficient $D$ of the motile fraction of a population can be estimated from measured bacterial trajectories, according to $D = (1/6)v^2(R + 4D_R)/(R + 2D_R)^2$ [61], where $v$ is the average velocity of motile cells (Fig. S7A), $R$ is the reorientation frequency (Fig. S7B), and $D_R = 0.035\text{rad}^2/\text{s}$ is the rotational diffusion coefficient [57]. Using the bacterial swimming trajectories quantified at each time point during starvation, we computed the diffusion coefficient of each strain over time (Fig. S7C). The average effective diffusion coefficient of all limokinetic strains decreased from $1.5 \pm 0.5 \cdot 10^{-6}$ cm$^2$s$^{-1}$ in nutrient-rich media to $0.8 \pm 0.4 \cdot 10^{-6}$ cm$^2$s$^{-1}$ during starvation ($t$-test: $p = 0.003$, Fig. S7C).

Starved cells would benefit from increasing the amount of space they explore in search for new resources, i.e., their diffusion coefficient. They can do so by either increasing their swimming velocity $v$, which is energetically costly, or by lowering their reorientation rate (i.e., decreasing $R$), which comes at no energetic cost and would thus be a preferable adaptation. However, tracking showed that, rather than decreasing, the average reorientation frequency increases during starvation (Fig. S7B). This suggests that there might be additional reasons for cells not to decrease their reorientation frequency. The possibility that bacteria under starvation adopt forms of random walk that enable more exploration of space, and thus more encounters, such as Lévy-flights [64, 25], cannot be ruled out from our experiments, but would require longer trajectories that are challenging to obtain.

#### 2.2 Limokinetic motility as a strategy for oligotrophic waters

75 % of the open ocean are oligotrophic [36], having background concentrations of dissolved organic matter (DOM) insufficient to sustain the growth of most copiotrophic bacteria [31]. A comparison of carbon concentrations and uptake rates show that DOM concentrations in oligotrophic waters are also typically insufficient to sustain swimming. For example, a concentration of serine of 100 nM would generate an energy flux of $\sim 10^5$ ATP/s (based on a measured bacterial uptake rate of $1 \cdot 10^5$ molecules/cell/s [69, 52], a $K_M$ of 1 µM [2], and an energy equivalent per serine molecule of 12 ATP [29]). Thus, background concentrations of amino acids of 100 nM [13] measured in coastal regions of the ocean are expected to be sufficient to fuel swimming with a velocity of 60 µm/s (requiring $\sim 9 \cdot 10^4$ ATP/s, as calculated using the equation and parameters presented in the main text), but this would severely reduce the energy flux available for cellular maintenance. Lower concentrations of nutrients are likely insufficient to fuel swimming, although some copiotrophs can express highly specific transporters under starvation with values of $K_M$ as low as 10 nM [18]. However, this is unlikely to generate a large energetic flux as models of high-affinity transporters show that there is a trade-off between affinity and the maximum uptake rate [44]. Therefore, in some coastal regions the background nutrient concentrations in the ocean may provide sufficient energy to swim, but in most regions of the ocean bacteria exposed to the background concentrations of nutrients will lack sufficient nutrients to fuel swimming.

Typical search times of motile marine bacteria for organic matter particles are expected to be on the order of hours to weeks [34]. In the areas with highest primary productivity (Beaufort Sea), the concentration of particles reaches $c$ = 50,000 particles/L (data from Tara oceans [47], integrating over all particle sizes from 20 µm to 1 mm), which corresponds to a typical distance of $L = 1.5$ mm between particles ($L \approx 0.553c^{-1/3}$ [9]). Using the relation $t = L^2/6D$ for the time $t$ required to diffuse a distance $L$, and assuming a diffusion coefficient of $D = 1 \cdot 10^{-6}$ cm$^2$s$^{-1}$, this corresponds to a search time $t$ of approximately 1 hour. In such environments, a small fraction of limostatic strains may be able to find a new particle before motility ceases, whereas limokinetic strains are expected to easily find a new particle. The discrepancy between the two strategies increases as the particle concentration decreases. In areas with very low productivity the particle concentration is typically very low. For example in the South Pacific Ocean, at $c = 0.4$ particles/ L, corresponding to a typical distance of $L = 74$ mm between particles, the average search time $t$ for motile cells is more than 3 months. Given our measurements of the cost of bacterial motility (62 % biomass reduction over one week), it seems unlikely that motility endurance extends for multiple months. However, assuming exponentially distributed search times [34], even in this extremely oligotrophic environment 10% of the motile bacteria will have encountered a particle within one week. This indicates that the limokinetic strategy may still be viable in extremely nutrient-poor regions of the ocean, even when the average search time significantly exceeds the average swimming endurance time of the population.

#### 3 Supplementary Figures

Figure S1: **Phylogenomic tree of all strains used in this study**. Isolates spanning 3 orders of gammaproteobacteria were used in this study, with high representation among the Vibrionaceae. The names of studied strains are colored according to their behavioral response to starvation: limokinetic strains in blue; limostatic strains in orange. The names of additional strains used to test the predictive ability of the classifier are shown in red ('strains to test model', see Fig. 5 and Main Text). Node values represent Shimodaira-Hasegawa (SH)–like test values with 10,000 resamplings (obtained using the -alrt option in IQTree2; [40]). Phylogenomic tree was constructed using PhyloPhlAn 3.0 [1]. The tree is rooted on *Pelagibacter ubique 1062* (SAR11) (NCBI accession: CP000084.1; not shown). Tree pruning,ordering and aesthetics were carried out using ETE3. Scale bar represents 0.01 nucleotide substitutions per site.

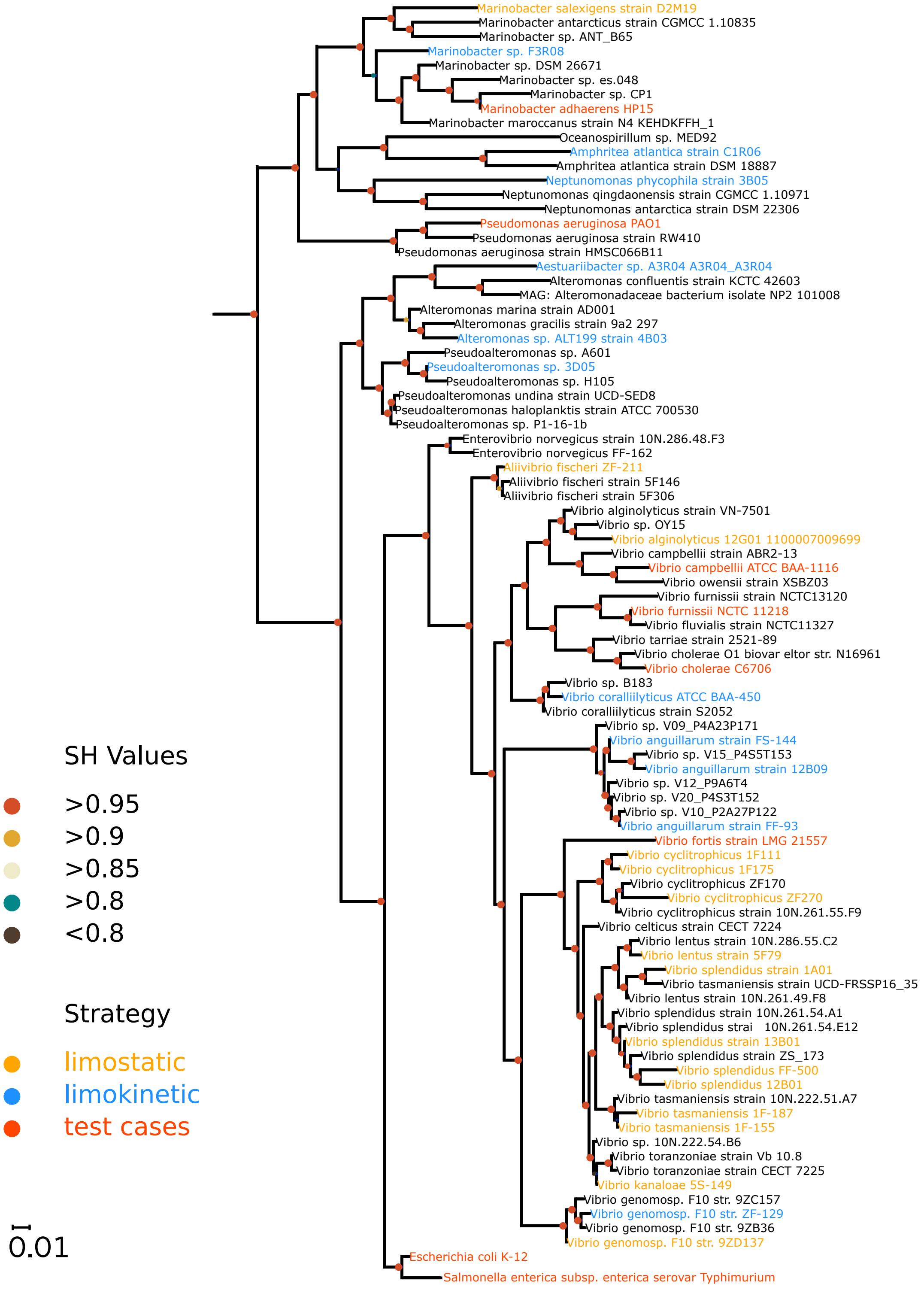

Figure S2: **Response of marine bacteria to starvation reveals a dichotomy in motility endurance.** **A**: Distribution of average swimming velocities in *V. splendidus* 1A01 (left) and *V. anguillarum* FS-144 (right) prior to starvation ('C+'; top) and at different times during carbon starvation (1 h to 47 h; bottom). Dashed gray lines mark the velocity of 12 μm/s, used to differentiate motile from non-motile cells (panel B). 'pdf': probability density function. Data originate from identical experiments as used to produce Fig 1A, for a different pair of strains from the same species. **B**: Cellular velocity distributions for 22 strains during growth in carbon-replete media, with a bin width of 3 μm/s. A threshold of 12 μm/s (dashed lines) was used to distinguish motile from non-motile cells in the population. **C**: The number of observations for each of the 28 strains during the starvation experiment (Fig 1). For each time bin, the number of independent experiments (performed on different days) is shown (number and heatmap), with the total number of videos (including replicates taken on the same day) indicated in brackets. **D**: Separation of limokinetic and limostatic strains based on a kernel-density estimate (KDE) on the logarithm of the time-averaged fraction of motile cells per strain, for starvation times ¿1 h. Results for two different bandwidths $\sigma$ are shown, $\sigma = 0.05$ (gray) and $\sigma = 0.10$ (black). A single local minimum (at 0.033) of the KDE indicates the fraction of motile cells that best separates two behavioral classes. **E**: Average swimming velocities of motile cells during growth for limokinetic and limostatic strains. The average velocities of the two behavioral classes differ significantly (Mann-Whitney U test (M.W.U.): $p = 0.02$). **F**: Fraction of motile cells during growth for limokinetic and limostatic strains. The average motile fractions of the two behavioral classes differ significantly (M.W.U.: $p = 0.005$).

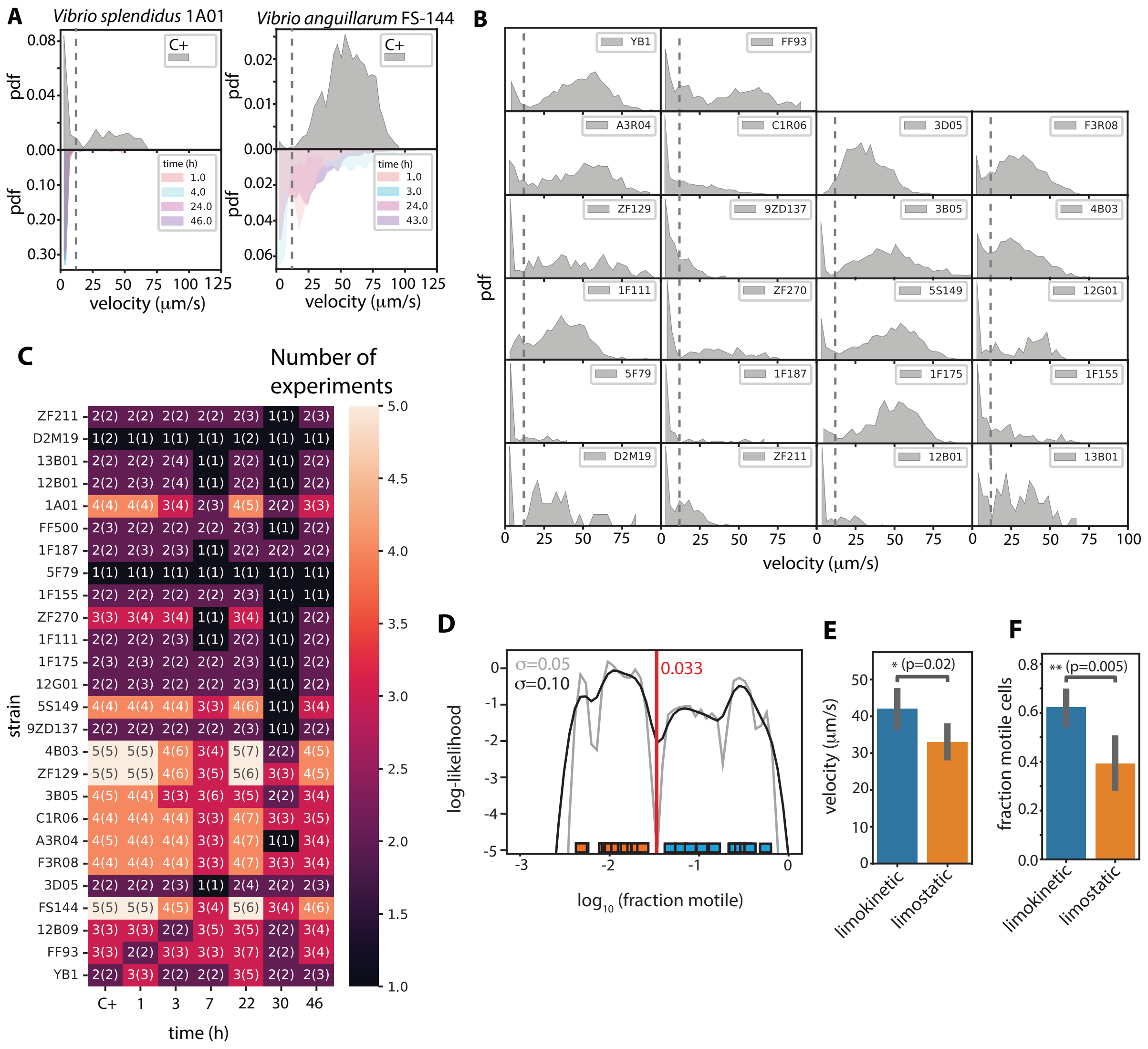

Figure S3: **The motility response to nutrient limitation is similar during stationary phase and after washing.** To quantify motility during stationary phase, we diluted cells (1/2000) from a culture grown overnight in 100% MB into 2% MB in artificial seawater. The concentration of MB was chosen to obtain a final concentration of cells that is low enough to remain compatible with tracking (OD $\approx 0.02$). A: Heat map showing the motile fraction for different strains at various time points in 2% in Marine Broth. The red vertical line indicates the approximate transition between growth and stationary phase, based on the plateau in the cell number in panel B. Strain names are shown in blue for limokinetic strains and orange for limostatic strains, based on their motility response after washing and transfer to carbon-depleted medium. B: Number of cells (measured as the average number of trajectories per frame; blue) and fraction of motile cells (red) as a function of time in 2 % MB. C: The time-averaged fraction of motile cells per strain in starvation medium and in stationary phase is highly correlated (Pearson's $\rho$ = 0.91, CI: 0.67–0.97). Blue indicates limokinetic strains and orange limostatic.

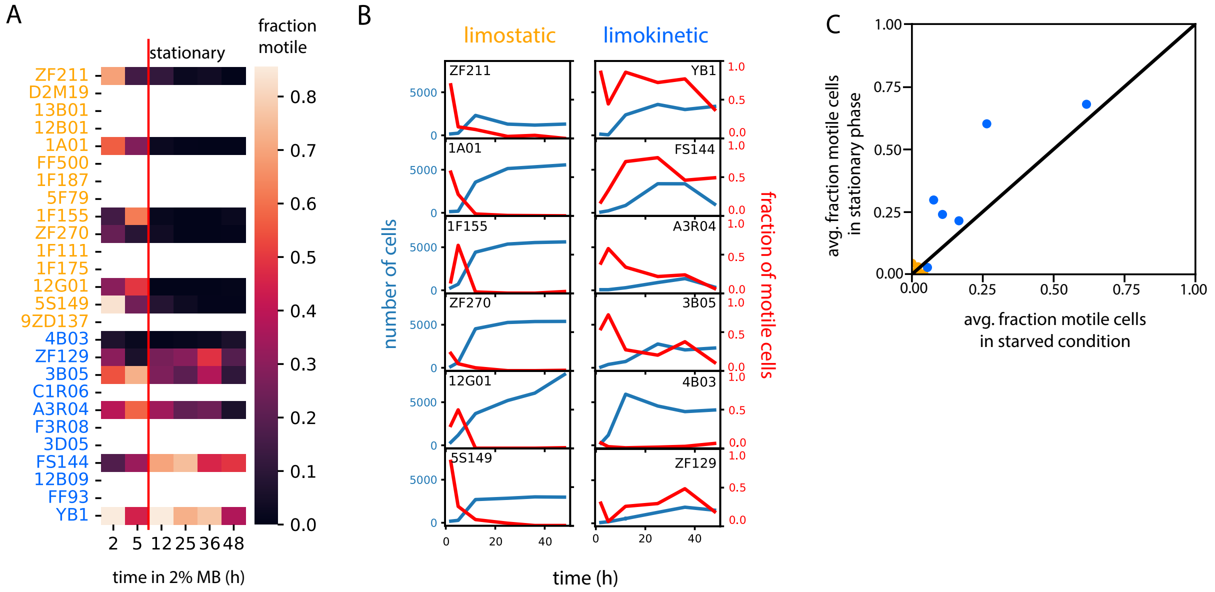

Figure S4: **Loss of flagellar filaments is a result of exposure to nutrient starvation, not mechanical stress.** A: The fraction of motile cells (top) and average velocity per cell (bottom) for cells growing in 50% Marine Broth ('before centrifugation', yellow), after washing in starvation buffer (purple), and after washing in fresh 50% Marine Broth (red), for seven strains (four limostatic strains labeled in orange and three limokinetic in blue). B: (top) Example image of 24 h-starved *Vibrio cyclitrophicus* ZF270 cells with an isolated flagellar filament. Scale bar 10 µm. (bottom) Image of a single isolated flagellar filament from a sample of 3 h-starved ZF270. Scale bar 2 µm. C: Distribution (shaded area) and KDE-estimate (lines) of filament length for flagella of all measured limokinetic (blue, $N = 189$) and limostatic (orange, $N = 388$) cells. Filaments measured by SEM in an experiment where the cells were grown in carbon-replete media and then starved for up to 24 h (Fig 2). Distributions are not significantly different (M.W.U: $p = 0.26$). D: Distribution (shaded area) and KDE-estimate (lines) of flagellar length for all cells during growth in carbon-replete media (yellow, $N = 268$) and during carbon starvation (purple, $N = 309$). Distributions are not significantly different (M.W.U: $p = 0.09$). Data originates from the same experiment as in panel C.

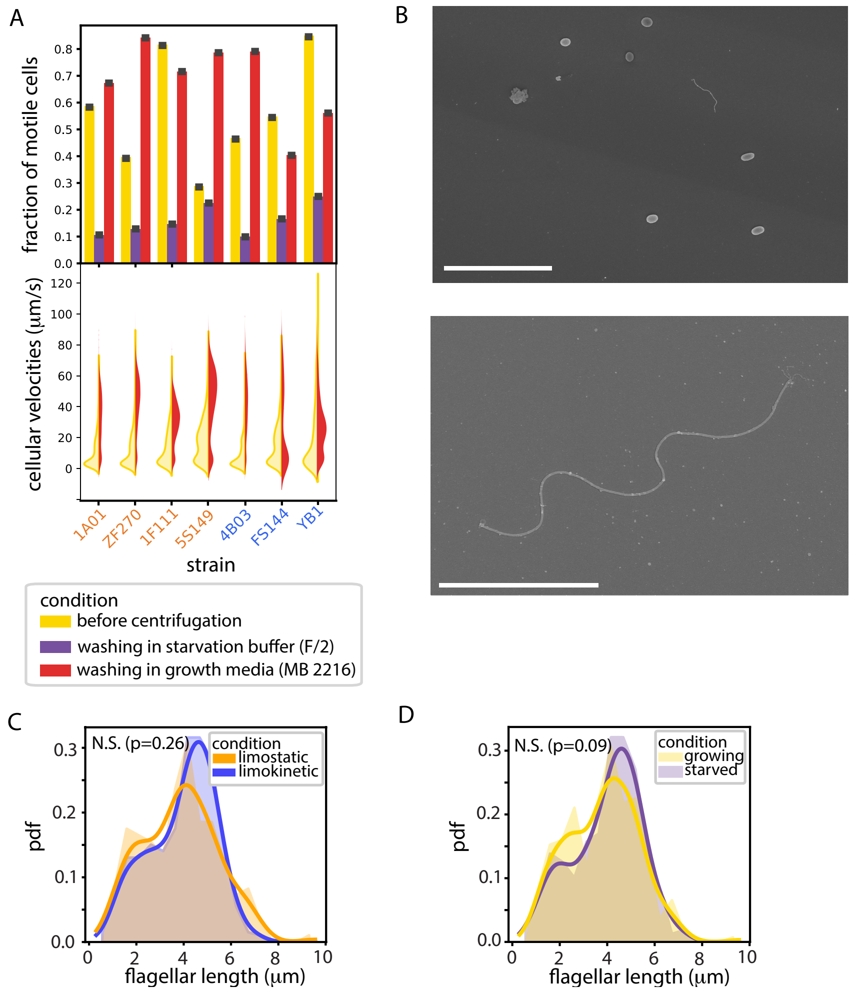

Figure S5: **Delayed motility recovery in limostatic compared to limokinetic strains.** Fraction of motile cells (A) and average velocities of motile cells (B) for limostatic (orange) and limokinetic (blue) strains after the addition of 1% Marine Broth to cultures starved for 24 h. Velocities are shown for strains in which the fraction of motile cells was greater than 10%.

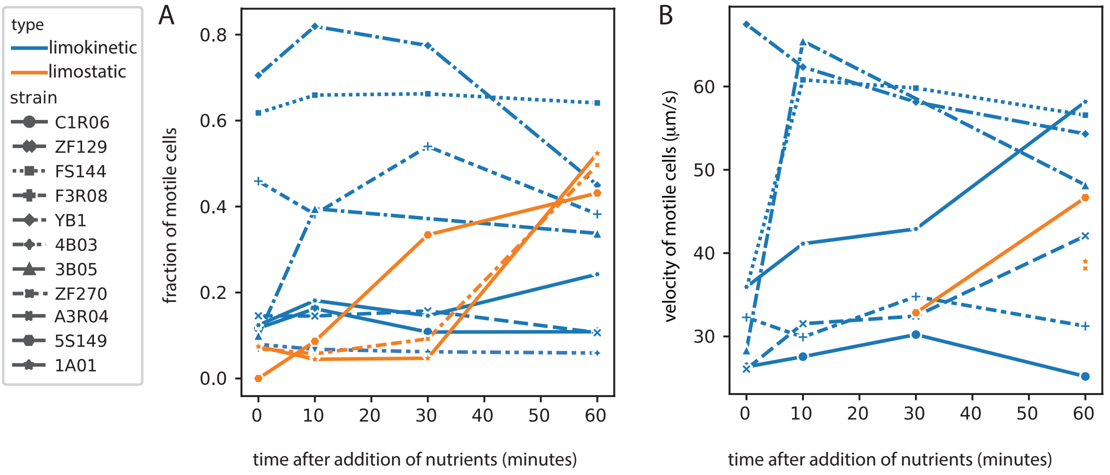

Figure S6: **Relation between motility during growth. A**: Relationship between specific growth rate (ln(2)/doubling time) and average speed of motile cells during growth in 50% Marine Broth for 23 strains. The correlation coefficient, $\rho_p = 0.02$ (dashed line), is shown with 95% confidence interval (shaded area). B: Relationship between specific growth rate and fraction of motile cells for 23 strains. The correlation coefficient, $\rho_p = 0.29$ (dashed line), is shown with 95% confidence interval (shaded area). **C**: Average growth rates during growth for limokinetic and limostatic strains are not significantly different (M.W.U.: $p = 0.09$).

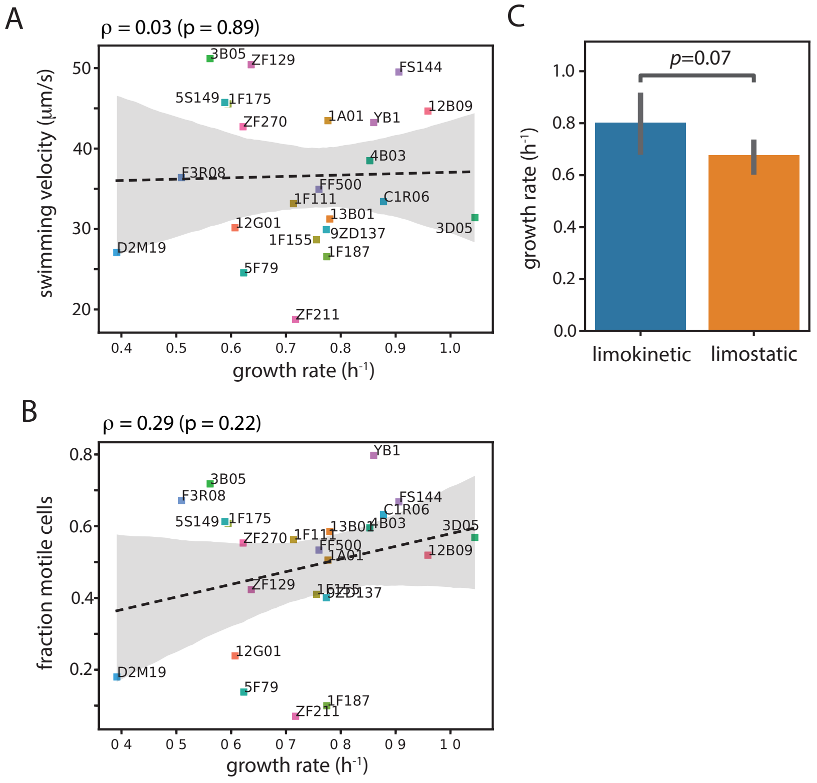

Figure S7: **Bacterial motility parameters during starvation. A**: Population-averaged swimming velocity of motile cells as a function of time for single strains (gray lines) and the average over all strains (blue line) with 95% confidence interval (shaded area). 'C+' denotes the condition prior to starvation. **B**: Population-averaged reorientation frequency as a function of time for single strains (gray lines) and the average over all strains (blue line) with 95% confidence interval (shaded area). 'C+' denotes the condition prior to starvation. To detect reorientations during swimming, cellular positions were processed with a second-order Savitzky–Golay filter [51] with a time window of 5 frames to compute the angle and velocity between frames. For each trajectory, reorientation events were identified as time points at which both 1) the absolute change in angle exceeded 25°, and 2) the velocity was lower than 75% of the average velocity of the trajectory. The minimal time between two reorientation events was limited to 2 frames (60–80 ms). The run time was defined as the time between detected reorientation events. The first run (from the start of the trajectory to the first event) and the last run (from the last detected event to trajectory length) were used as lower bound estimates of the run time. The reorientation frequency per cell was calculated as the inverse of the mean average run time per cell. To prevent detection of spurious events in slowly moving cells, the analysis was only applied to trajectories with a minimum length of 30 frames, and a minimum velocity of 12 μm/s based on average-filtered positional data with a time window of 9 frames. **C**: The effective diffusion coefficient of all strains (colored lines) as a function of time. The average diffusion coefficient for the motile fraction of the population was computed as $D = (1/6)v^2(R + 4D_R)/(R + 2D_R)^2$ [61], where $v$ is the average velocity of motile cells (panel A), $R$ is the reorientation frequency (panel B), and $D_R = 0.035\text{rad}^2/\text{s}$ the rotational diffusion coefficient [57]. 'C+' denotes the condition prior to starvation

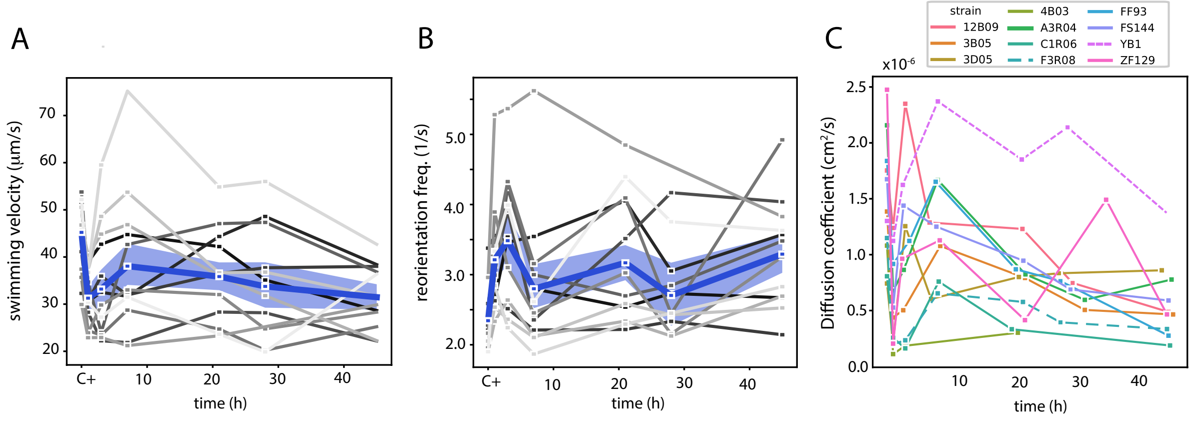

Figure S8: **Cell counts and motility in a week-long starvation experiment.** **A:** Fraction of motile cells of three limokinetic strains as a function of starvation time for times >48 h. **B:** Number of cells as a function of time, as estimated from the number of trajectories per frame, normalized to the estimate at the first time point after washing. Error bars 95% CI. **C:**: Viable cell concentration as a function of time, measured by colony counting on Marine Broth (1.5% agar) plates. Only plates with 20–350 colonies were included in the analysis. Error bars denote the standard deviation of 2–3 plates per condition. **D:** KDE-distributions of cell length from SEM as a function of starvation time. Note that actual cell lengths may deviate from SEM estimates due to desiccation during sample preparation, but relative changes are expected to be conserved. The average cell size decreases by 31%, consistent with reductive divisions taking place after the onset of starvation. **E**: Fraction of cells with a compromised membrane as a function of time. The number of dead cells was estimated using SYTOX Green staining and the total number of cells by SYBR Green staining (Fig 3C).

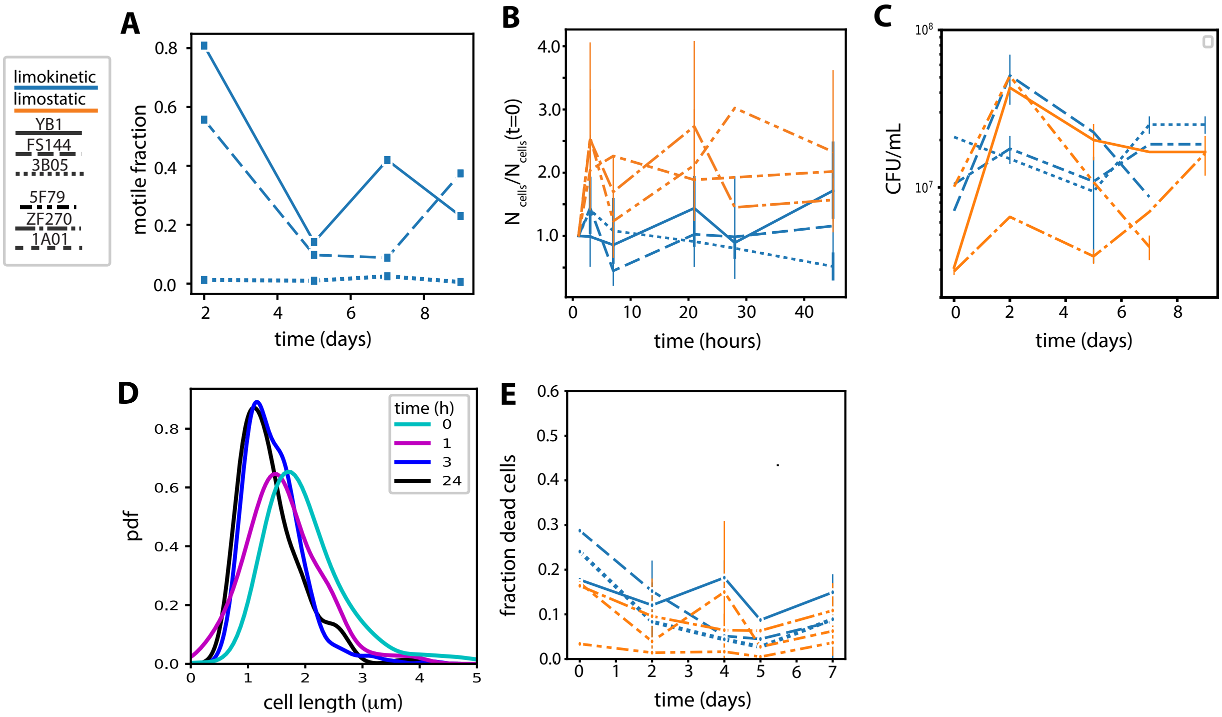

Figure S9: **Alternative energy sources. A:** Limokinetic cells are also motile in the dark. Average swimming velocity (top) and motile fraction (bottom) of cells during starvation ($t$ ¿ 24 h) in the dark (dark gray, single replicate) and with normal light exposure (pale gray, see Fig. 1). To test the effect of light on swimming, *Vibrio anguillarum* FS144 and *Vibrio coralliilyticus* YB1 cultures were grown and starved following the standard protocol (Materials and Methods) but then kept in culture tubes wrapped in aluminium foil. Microscopy samples were prepared in the dark and cell motility was quantified immediately upon placing them on the microscope. Without covering the tubes, the cells experienced a diel cycle (approx. 16 h of light per day) with the starvation process starting in the afternoon. Error bars indicate one S.D. **B:** The motile fraction of cells, for starvation times $>$ 24 h, as a function of cell concentration for four limokinetic strains (colored points). For each strain, the cell concentrations were obtained by diluting the same culture, where 1 indicates the standard dilution in our starvation protocol, corresponding to $\sim 10^7$ mL$^{-1}$. Solid lines represent linear fits to the data, with the slope $\beta$, residuals $R^2$ and associated probability $p$ indicated in the figure legend. The motile fraction is not negatively correlated with the cell concentration, which excludes there being a large influence on motility of residual nutrients in the medium.

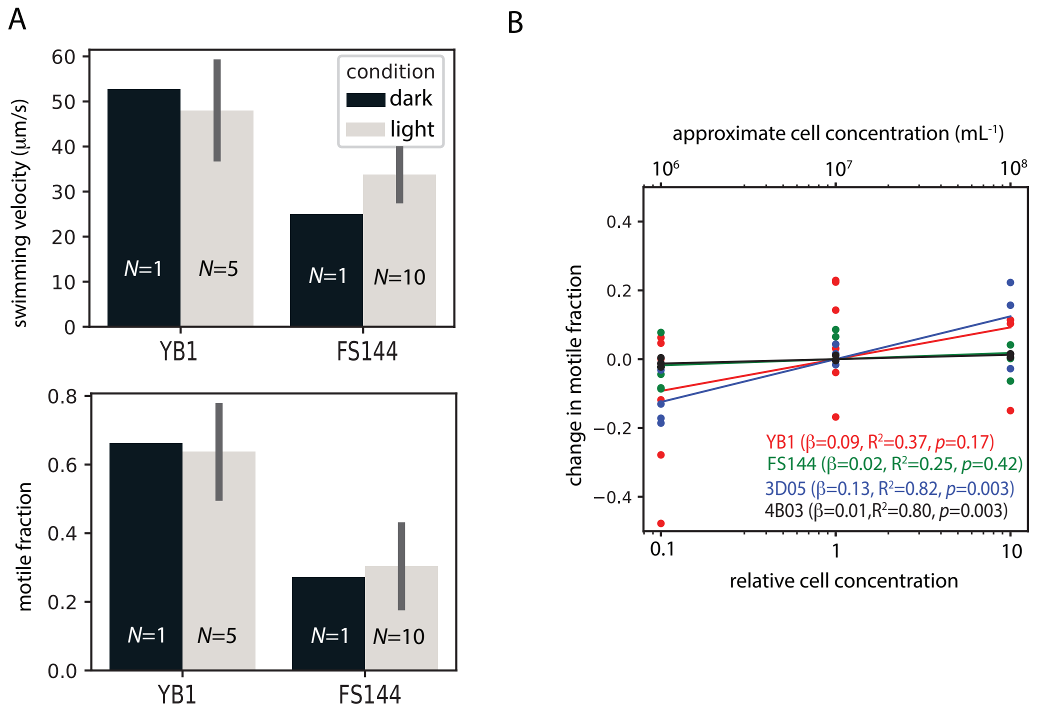

Figure S10: **Polyphosphate granules in *Alteromonas sp.* 4B03** A: Representative images of cells stained with DAPI, which stains both DNA (blue) and polyphosphate (yellow/green dots, panel A), for four limokinetic strains and two limostatic strains when starved for 3h. PolyP granules were only observed in *Alteromonas sp. 4B03*. B: Fraction of cells of *Alteromonas sp.* 4B03 containing at least 1 polyphosphate granule per cell, for cells stained during growth (0 h) and cells in carbon-limited medium (3 h). In the carbon-limited medium, phosphate is in excess (Materials and Methods). C: Average number of polyphosphate granules per cell under the same two conditions. $N$ indicates the number of cells per condition.

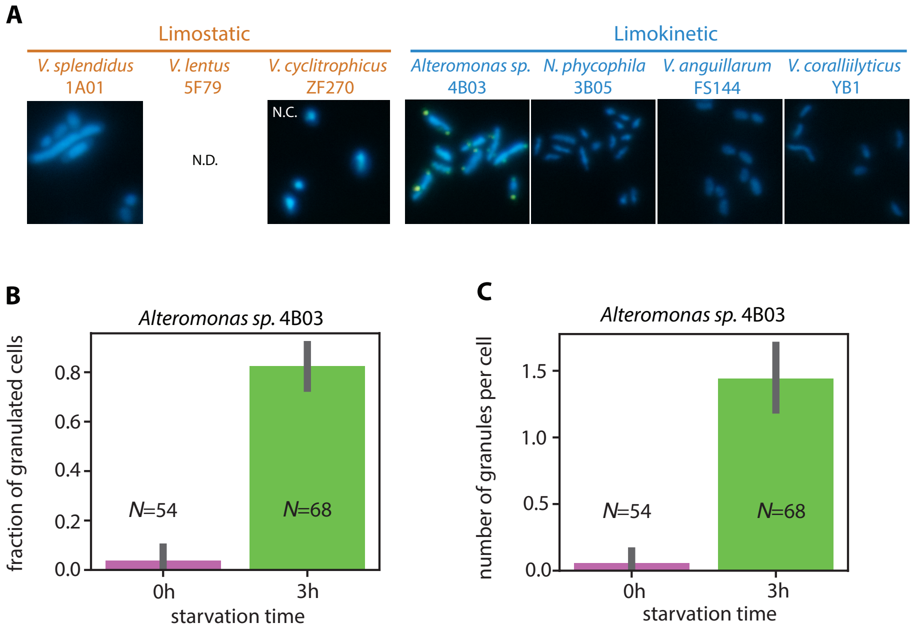

Figure S11: Recursive feature elimination (RFE) for the training set of our limokinetic Bayesian classifier (green) compared to the mean (red) and mean plus standard deviation (orange, dashed) of 100 trials in which class (limostatic/limokinetic) was randomly assigned. RFE was run for combinations of 2, 4, 8, 16, 32, 64, 128, 256, 512, and 1000 features.

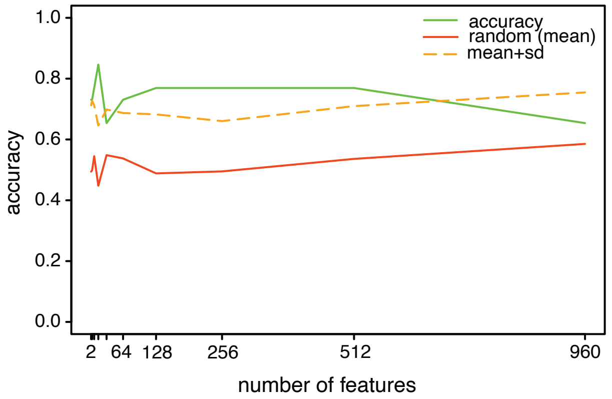

Figure S12: **Influence of oxidative stress on growth, motility, and lag time.** A: Growth rate as a function of the added external hydrogen peroxide concentration [$H_2O_2$] for three limokinetic (blue) and three limostatic strains (orange). Growth rates were normalized to the growth rate of each strain measured without the addition of hydrogen peroxide. B: Average swimming velocity as a function of [$H_2O_2$] for three limokinetic (blue) and two limostatic strains (orange). Velocities were normalized to the velocity of each strain without the addition of hydrogen peroxide. C: Lag time as a function of the added [$H_2O_2$] for three limokinetic (blue) and three limostatic strains (orange). Cultures were starved for 24 h and lag time was measured after adding Marine Broth (final concentration 50%). Lag time was defined as the time until the culture reached an OD of 0.05. Values were normalized by the lag time of each strain without the addition of hydrogen peroxide. The average lag time at [$H_2O_2$] = 10 µm was not significantly different ($t$-test, $p = 0.24$) between limostatic (1.36) and limokinetic (1.09) strains.

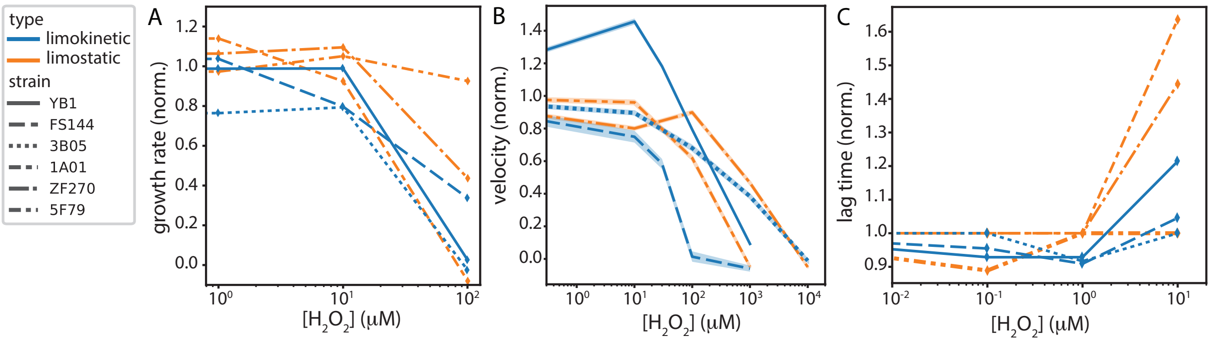

Figure S13: Prevalence of orthologous groups (OG) associated with a limostatic response for both limokinetic and limostatic strains, as obtained by RFE (Methods) and clustered into 15 functional categories. OG with significance $p < 0.05$ from a regression analysis that includes phylogeny (Materials and Methods) are marked with *. Circles indicate the gene copy number of each OG (size) and the probability of the association with the limostatic response (color). 'Prediction' (bottom row) is the predicted class of each strain by the classifier.

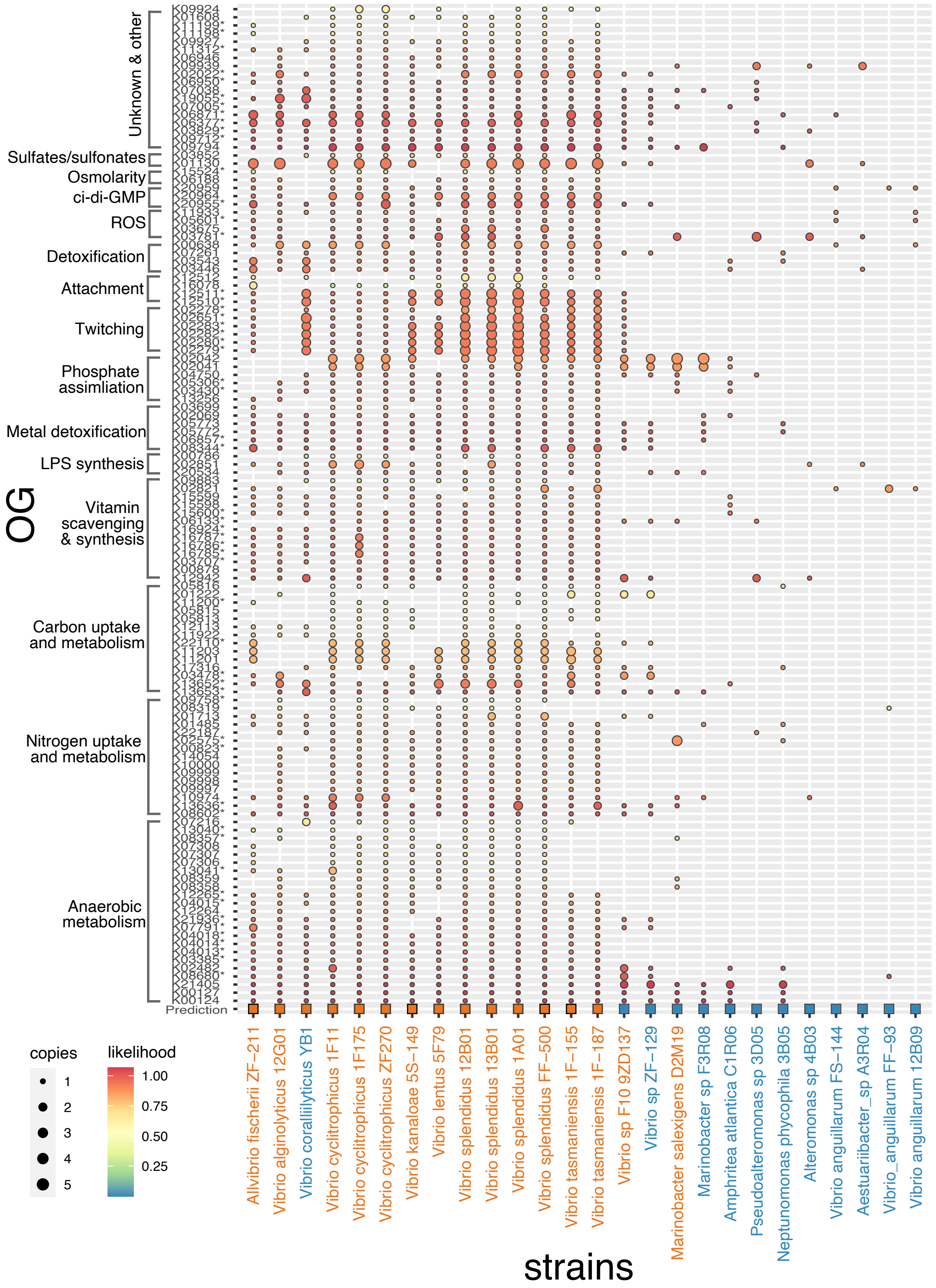

Figure S14: A: Testing of classifier prediction in other marine strains not used for initial training. Lines show the observed motile fraction as a function of starvation time under the same conditions as in Fig. 1, providing experimentally-determined classification (colors). 'G' denotes the condition before starvation. B: Testing of classifier prediction in enteric bacteria. Shown are the measured motile fraction of *E. coli* RP437 grey) and *Salmonella typhimurium* LT2 (black) and fits to a single exponential decay (dashed lines). For these strains, the standard starvation protocol was adapted by replacing Marine Broth with Tryptone broth (10 g/L Bacto tryptone and 5 g/L NaCl), and as starvation buffer adapted motility buffer [55] was used without potential energy sources Lactic acid and methionine (10 mM potassium phosphate at pH 7.0, 0.1 mM EDTA). Each experiment was replicated twice. The behavior of the enteric bacteria deviates from the strong dichotomy found in marine strains: their motile fraction steadily decreases with starvation time, representing limostatic behavior, but with a timescale much longer than observed in marine strains (orange). Their motile fraction is much lower compared to the average of all limokinetic marine strains (blue). Shaded areas and error bars represent to 95 % CI. Dashed lines indicate an exponential fit of the motile fraction as a function of starvation time (excluding the datapoint for growth). Inset: motility loss timescale for limostatic strains, obtained from single exponential fits to the motile fraction during starvation.

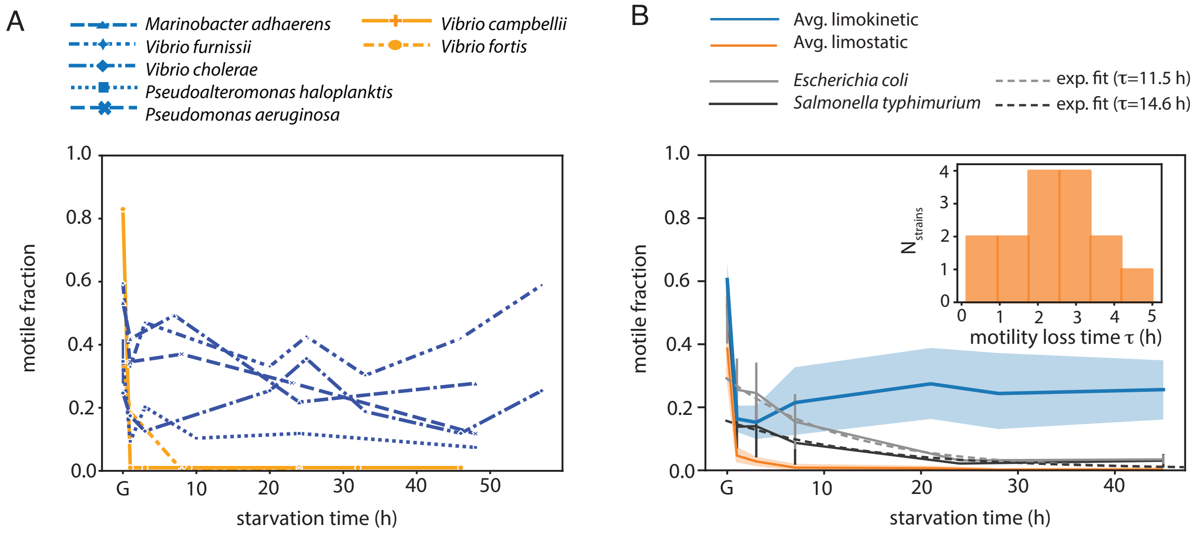

Figure S15: Predicted fraction of limokinetic (blue) and limostatic (orange) taxa for different ocean sampling time points (circles), normalized by all gamma proteobacterial taxa. Prediction is based on the limostatic and limokinetic classifier (Figs. 5 and S13), including only the taxa where the both classifiers have identical prediction (e.g. excluding ambigious predictions). The line represents a Loess smoothed average.

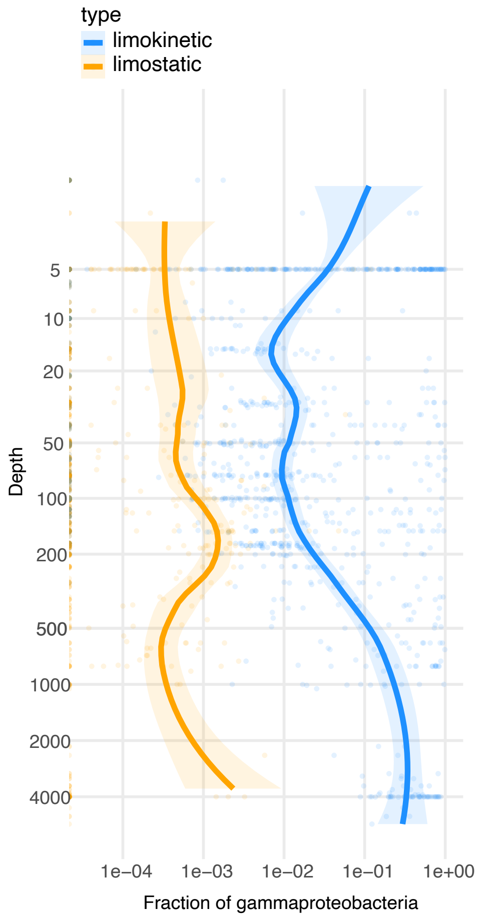

#### 4 Supplementary Tables

Table 1: Motility parameters for each strain during growth in carbon-replete medium (g) and during carbon starvation* (s). Fraction of motile cells $f$, average swimming velocity of motile cells $\langle v \rangle$ (µm/s), reorientation frequency $R$ (1/s).

| Strain | Species | accession | source | Category | $f_g$ | $f_s$ | $\langle v \rangle_g$ | $\langle v \rangle_s$ | $R_g$ | $R_s$ |
|---|---|---|---|---|---|---|---|---|---|---|
| ZF211 | *Aliivibrio fischeri* | GCF_000287175.2 | [24] | Limostatic | 0.07 | 0.000 | 19 | – | 5.5 | – |
| D2M19 | *Marinobacter salexigens* | GCF_018860765.1 | [14] | Limostatic | 0.18 | 0.000 | 27 | – | 4.1 | – |
| 13B01 | *Vibrio splendidus* | GCF_001691275.1 | [24] | Limostatic | 0.59 | 0.005 | 31 | – | 1.8 | – |
| 12B01 | *Vibrio splendidus* | GCF_000152765.1 | [24] | Limostatic | 0.40 | 0.012 | 37 | – | 2.3 | – |
| 1A01 | *Vibrio splendidus* | GCF_002700025.1 | [12] | Limostatic | 0.51 | 0.001 | 43 | – | 1.6 | – |
| FF-500 | *Vibrio splendidus* | GCF_000272265.2 | [24] | Limostatic | 0.49 | 0.007 | 39 | – | 1.1 | – |
| 5F79 | *Vibrio lentus* | GCF_001691195.1 | [24] | Limostatic | 0.14 | 0.015 | 25 | – | 2.4 | – |
| 1F187 | *Vibrio tasmaniensis* | GCF_000272405.2 | [24] | Limostatic | 0.10 | 0.012 | 27 | – | 3.7 | – |
| 1F155 | *Vibrio tasmaniensis* | GCF_000272385.2 | [24] | Limostatic | 0.41 | 0.020 | 33 | – | 2.2 | – |
| ZF270 | *Vibrio cyclitrophicus* | GCF_000256465.2 | [24] | Limostatic | 0.55 | 0.014 | 43 | – | 2.5 | – |
| 1F111 | *Vibrio cyclitrophicus* | GCF_000247005.2 | [24] | Limostatic | 0.56 | 0.01 | 33 | – | 1.5 | – |
| 1F175 | *Vibrio cyclitrophicus* | GCF_000256135.2 | [24] | Limostatic | 0.61 | 0.01 | 46 | – | 1.6 | – |
| 12G01 | *Vibrio alginolyticus* | GCF_000153505.1 | [24] | Limostatic | 0.24 | 0.01 | 30 | – | 2.5 | – |
| 5S149 | *Vibrio kanaloae* | GCF_000272165.2 | [24] | Limostatic | 0.61 | 0.02 | 46 | – | 2.0 | – |
| 9ZD137 | *Vibrio sp. F10* | GCF_000287015.2 | [24] | Limostatic | 0.40 | 0.02 | 30 | – | 2.5 | – |
| 4B03 | *Alteromonas sp.* | GCF_018619335.1 | [12] | Limokinetic | 0.60 | 0.048 | 38 | 22 | 2.6 | 3.5 |
| ZF129 | *Vibrio sp.* | GCF_000287055.2 | [24] | Limokinetic | 0.42 | 0.062 | 50 | 26 | 1.9 | 3.6 |
| 3B05 | *Neptunomonas phycophila* | GCF_018619275.1 | [12] | Limokinetic | 0.72 | 0.080 | 51 | 36 | 3.4 | 3.6 |
| C1RO6 | *Amphritea atlantica* | GCF_018860855.1 | [14] | Limokinetic | 0.63 | 0.110 | 33 | 21 | 2.4 | 3.9 |
| A3R04 | *Aestuariibacter sp.* | GCF_018860805.1 | [14] | Limokinetic | 0.57 | 0.150 | 52 | 34 | 2.0 | 3.2 |
| F3R08 | *Marinobacter sp.* | GCF_018860425.1 | [14] | Limokinetic | 0.67 | 0.25 | 36 | 27 | 2.2 | 2.9 |
| 3D05 | *Pseudoalteromonas sp.* | GCF_002723455.1 | [12] | Limokinetic | 0.57 | 0.33 | 31 | 35 | 2.3 | 2.2 |
| FS144 | *Vibrio anguillarum* | GCF_000287115.2 | [24] | Limokinetic | 0.67 | 0.25 | 50 | 38 | 2.1 | 2.4 |
| 12B09 | *Vibrio anguillarum* | GCF_000287135.2 | [24] | Limokinetic | 0.52 | 0.290 | 45 | 37 | 2.2 | 2.8 |
| FF93 | *Vibrio anguillarum* | GCF_000287095.2 | [24] | Limokinetic | 0.49 | 0.370 | 43 | 33 | 1.9 | 2.4 |
| YB1 | *Vibrio coralliilyticus* | GCF_000176135.1 | [67] | Limokinetic | 0.80 | 0.550 | 43 | 53 | 2.5 | 2.4 |
| | average ± std ** | | | All | 0.48±0.19 | 0.10±0.15 | 37±9 | – | 2.44 ±0.87 | – |
| | average ± std | | | Limostatic | 0.39±0.20 | 0.01±0.01 | 33±8 | – | 2.54 ±1.08 | – |
| | average ± std | | | Limokinetic | 0.61±0.11 | 0.23±0.16 | 43±7 | 33±9 | 2.30 ±0.42 | 3.0±0.6 |

*:for all averages during starvation, all times >1 h are included.
**:average and standard deviation of the average number per strain.

Table 2: Presence (+) or absence (+) of the main storage compound synthesis and catabolysis genes for each species, based on annotated genomes (accession numbers in Table S1) and annotations according to BioCyc [30]. Presence/absence are the same for all strains of the same species unless noted otherwise.

| | | PHB | | | PolyP | | glycogen | |
|---|---|---|---|---|---|---|---|---|
| Species | Category | *phaB* | *phaC* | *phaZ* | *ppk1* | *ppk2* | *glgC* | *glgP* |
| *Aliivibrio fischeri* | Limostatic | - | - | - | - | - | + | + |
| *Marinobacter salexigens* | Limostatic | + | + | - | + | - | - | - |
| *Vibrio splendidus* | Limostatic | +* | +* | - | + | - | + | + |
| *Vibrio lentus* | Limostatic | + | + | - | + | - | + | + |
| *Vibrio tasmaniensis* | Limostatic | - | + | - | + | - | + | + |
| *Vibrio cyclitrophicus* | Limostatic | + | + | - | +** | - | + | + |
| *Vibrio alginolyticus* | Limostatic | + | + | - | + | + | + | + |
| *Vibrio kanaloae* | Limostatic | + | + | - | + | - | + | + |
| *Vibrio sp. F10* | Limostatic | + | + | - | - | - | + | + |
| *Alteromonas sp. 4B03* | Limokinetic | - | - | - | + | + | + | + |
| *Vibrio sp. ZF-129* | Limokinetic | + | - | - | - | - | + | + |
| *Neptunomonas phycophila* | Limokinetic | + | + | + | + | + | - | - |
| *Amphritea atlantica* | Limokinetic | + | + | - | + | - | - | - |
| *Aestuariibacter sp.* | Limokinetic | + | + | + | + | + | + | + |
| *Marinobacter sp.* | Limokinetic | + | + | + | + | + | + | + |
| *Pseudoalteromonas sp.* | Limokinetic | - | - | - | + | + | + | + |
| *Vibrio anguillarum* | Limokinetic | - | - | - | + | - | + | + |
| *Vibrio coralliilyticus* | Limokinetic | + | + | - | + | + | + | + |

* *phaC* absent in 12B01 only, *phaB* absent in strains FF-500 and 13B01.
** *ppk1* absent in strain 1F111

Table 3: Results of regression analysis with and without phylogeny for limokinetic classifer.

| ModelName_R | Feature | Model | Intercept | Intercept p-value | Slope | Slope p-value |
|---|---|---|---|---|---|---|
| glm1 | K07666 | logistic | -19.57 | 0.995 | 20.87 | 0.995 |
| phyloglm1 | K07666 | plogistic | -17.091 | 0.9908 | 18.157 | 0.9902 |
| phylolm1 | K07666 | linear | 1.2698E-08 | 1 | 0.7825 | 3.9430E-06 |
| glm2 | K03782 | logistic | -2.565 | 0.01345 | 4.174 | 0.00127 |
| phyloglm2 | K03782 | plogistic | -1.38293 | 0.14323 | 2.81141 | 0.01194 |
| glm3 | K01507 | logistic | -2.485 | 0.01697 | 3.689 | 0.00274 |
| phyloglm3 | K01507 | plogistic | -1.07273 | 0.27514 | 2.4017 | 0.02938 |
| glm4 | K03606 | logistic | -2.485 | 0.01697 | 3.689 | 0.00274 |
| phyloglm4 | K03606 | plogistic | -1.0048 | 0.31616 | 2.5083 | 0.02024 |
| glm5 | K16554 | logistic | -2.485 | 0.01697 | 3.689 | 0.00274 |
| phyloglm5 | K16554 | plogistic | -1.0048 | 0.31616 | 2.5083 | 0.02024 |
| glm6 | K01690 | logistic | -1.8718 | 0.01373 | 3.3759 | 0.00195 |
| phyloglm6 | K01690 | plogistic | -1.16387 | 0.1546 | 2.42296 | 0.0268 |
| glm7 | K02302 | logistic | -1.8718 | 0.01373 | 3.3759 | 0.00195 |
| phyloglm7 | K02302 | plogistic | -1.082 | 0.1919 | 2.36317 | 0.03151 |
| glm8 | K04047 | logistic | -1.8718 | 0.01373 | 3.3759 | 0.00195 |
| phyloglm8 | K04047 | plogistic | -1.082 | 0.1919 | 2.36317 | 0.03151 |
| glm9 | K07645 | logistic | -1.8718 | 0.01373 | 3.3759 | 0.00195 |
| phyloglm9 | K07645 | plogistic | -0.5467 | 0.56215 | 2.38287 | 0.02792 |
| glm10 | K20977 | logistic | -1.8718 | 0.01373 | 3.3759 | 0.00195 |
| phyloglm10 | K20977 | plogistic | -0.5943 | 0.51001 | 2.28979 | 0.03643 |
| glm11 | K20978 | logistic | -1.8718 | 0.01373 | 3.3759 | 0.00195 |
| phyloglm11 | K20978 | plogistic | -0.5943 | 0.51001 | 2.28979 | 0.03643 |
| glm12 | K03411 | logistic | -1.5404 | 0.01547 | 3.6199 | 0.00342 |
| phyloglm12 | K03411 | plogistic | -0.38988 | 0.67737 | 2.65047 | 0.03924 |
| glm13 | K20920 | logistic | -1.7918 | 0.01898 | 2.8904 | 0.00436 |
| phyloglm13 | K20920 | plogistic | -0.41944 | 0.65052 | 1.87596 | 0.07197 |
| glm14 | K20988 | logistic | -1.7918 | 0.01898 | 2.8904 | 0.00436 |
| phyloglm14 | K20988 | plogistic | -0.41944 | 0.65052 | 1.87596 | 0.07197 |
| glm15 | K09797 | logistic | -1.4663 | 0.02206 | 2.8526 | 0.00505 |
| phyloglm15 | K09797 | plogistic | -0.54034 | 0.4978 | 1.74001 | 0.1058 |
| glm16 | K00140 | logistic | -1.4663 | 0.02206 | 2.8526 | 0.00505 |
| phyloglm16 | K00140 | plogistic | -0.74432 | 0.37128 | 1.84095 | 0.08248 |
| glm17 | K00523 | logistic | -1.2528 | 0.02713 | 3.1987 | 0.00821 |
| phyloglm17 | K00523 | plogistic | -0.57915 | 0.48964 | 2.14018 | 0.07855 |
| glm18 | K00641 | logistic | -1.2528 | 0.02713 | 3.1987 | 0.00821 |
| phyloglm18 | K00641 | plogistic | -0.71518 | 0.37987 | 2.25099 | 0.06539 |
| glm19 | K03410 | logistic | -1.0986 | 0.0334 | 19.6647 | 0.9941 |
| phyloglm19 | K03410 | plogistic | -0.22272 | 0.752 | 3.09185 | 0.109 |
| phylolm19 | K03410 | linear | 0.64855 | 0.07652 | 0.72675 | 0.03339 |
| glm20 | K12972 | logistic | -1.0986 | 0.0334 | 19.6647 | 0.9941 |
| phyloglm20 | K12972 | plogistic | -0.64003 | 0.39286 | 3.13663 | 0.06656 |
| phylolm20 | K12972 | linear | 0.13085 | 0.7805 | 0.88191 | 0.051 |
| glm21 | K02106 | logistic | -1.0986 | 0.0334 | 19.6647 | 0.9941 |
| phyloglm21 | K02106 | plogistic | -0.64003 | 0.39286 | 3.13663 | 0.06656 |
| phylolm21 | K02106 | linear | 0.13085 | 0.7805 | 0.88191 | 0.051 |
| glm22 | K03521 | logistic | -1.1787 | 0.0393 | 2.4314 | 0.0135 |
| phyloglm22 | K03521 | plogistic | -0.53876 | 0.5269 | 1.58075 | 0.1576 |

Table 4: Presence (+) or absence (-) of vps-related genes based on annotated genomes (accession numbers in Table S1) of all Vibrionaceae, including the (top) 11 species used to train the classifiers and the (bottom) four additional species used for prediction. Presence of *vps* genes refers to the presence of at least one of the *vpsN, vpsO, vpsM* genes. For *rbmC* genes, the predicted presence based on available genomes was corrected using the more specific annotation of Ref. [23].

| Species | Category | *vps* | *rbmC* |
|---|---|---|---|
| *Aliivibrio fischeri* | Limostatic | + | - |
| *Vibrio splendidus* | Limostatic | - | - |
| *Vibrio lentus* | Limostatic | - | - |
| *Vibrio tasmaniensis* | Limostatic | - | - |
| *Vibrio cyclitrophicus* | Limostatic | - | - |
| *Vibrio alginolyticus* | Limostatic | + | - |
| *Vibrio kanaloae* | Limostatic | - | - |
| *Vibrio sp. F10* | Limostatic | + | - |
| *Vibrio sp. ZF-129* | Limokinetic | + | - |
| *Vibrio anguillarum* | Limokinetic | + | +* |
| *Vibrio coralliilyticus* | Limokinetic | + | + |
| *Vibrio fortis* | Limostatic | - | - |
| *Vibrio campbellii* | Limostatic | + | - |
| *Vibrio furnissii* | Limokinetic | + | + |
| *Vibrio cholerae* | Limokinetic | + | + |

*Except strain FS144.

### 5 Supplementaty Videos

**Supplementary Video 1**: Example movie of tracked cells of *Vibrio splendidus* FF-500 in carbon-replete medium (50% MB). Unless noted otherwise, all supplementary videos are recorded at 25 fps and shown in real time, using 20X magnification and covering an area of 0.65 mm by 0.65 mm. Any drift visible in the videos was subtracted before trajectory analysis (Materials and Methods). The cellular velocity distribution corresponding to this video is shown in Fig. 1B (left, grey)

**Supplementary Video 2**: Example movie of tracked cells of *Vibrio anguillarum* 12B09 in carbon-replete medium (50% Marine Broth). The cellular velocity distribution corresponding to this video is shown in Fig. 1B (right, grey).

**Supplementary Video 3**: Example movie of tracked cells of *Vibrio splendidus* FF-500 after 23 h starvation. The cellular velocity distribution corresponding to this video is shown in Fig. 1B (left, purple)

**Supplementary Video 4**:Example movie of tracked cells of *Vibrio anguillarum* 12B09 after 24 h starvation. The cellular velocity distribution corresponding to this video is shown in Fig. 1B (right, purple)

**Supplementary Video 5**:Composite video of 9 strains (1 per species) without motility after 22–28 h of starvation. Video is shown with twice the acquisition speed and each panel covers an area of approximately 0.3 by 0.3 mm. From left to right and top to bottom: 1F155, 1F175, 5F79; 5S149, 9ZD137, 12G01; D2M19, FF500, ZF211.

**Supplementary Video 6**: Composite video of 9 strains (1 per species) with motility 22–28 h after starvation. Video is shown with twice the acquisition speed. From left to right and top to bottom: 12B09, F3R08, ZF129; YB1, 4B03, 3B05; A3R04, C1R06, 3D05.

**Supplementary Video 7**: Example movie of tracked cells of *Aestuariibacter* sp. A3R04 in carbon-replete medium (50% MB).

**Supplementary Video 8**: Example movie of tracked cells of *Vibrio lentus* 5F79 after 4 h starvation, where the population has lost motility.

**Supplementary Video 9**: Example movie of tracked cells of *Vibrio kanaloae* 5S-149 after 4 h starvation, with 2 motile cells in an otherwise non-motile population.

**Supplementary Video 10**: Example movie of tracked cells of *Vibrio coralliilyticus* YB1 after 8 h starvation, showing a high fraction of motile cells.

### References

[1] Francesco Asnicar, Andrew Maltez Thomas, Francesco Beghini, Claudia Mengoni, Serena Manara, Paolo Manghi, Qiyun Zhu, Mattia Bolzan, Fabio Cumbo, Uyen May, Jon G. Sanders, Moreno Zolfo, Evguenia Kopylova, Edoardo Pasolli, Rob Knight, Siavash Mirarab, Curtis Huttenhower, and Nicola Segata. Precise phylogenetic analysis of microbial isolates and genomes from metagenomes using PhyloPhlAn 3.0. *Nature Communications*, 11(1):2500, May 2020.

[2] B. Ayo, M. Unanue, I. Azúa, G. Gorsky, C. Turley, and J. Iriberri. Kinetics of glucose and amino acid uptake by attached and free-living marine bacteria in oligotrophic waters. *Marine Biology*, 138(5):1071–1076, May 2001.

[3] Federico Baltar, Thomas Reinthaler, Gerhard J. Herndl, and Jarone Pinhassi. Major Effect of Hydrogen Peroxide on Bacterioplankton Metabolism in the Northeast Atlantic. *PLoS ONE*, 8(4):e61051, April 2013.

[4] Erin M. Bertrand, Dawn M. Moran, Matthew R. McIlvin, Jeffrey M. Hoffman, Andrew E. Allen, and Mak A. Saito. Methionine synthase interreplacement in diatom cultures and communities: Implications for the persistence of B $_{12}$ use by eukaryotic phytoplankton. *Limnology and Oceanography*, 58(4):1431–1450, July 2013.

[5] Amandine Buffet, Eduardo P C Rocha, and Olaya Rendueles. Nutrient conditions are primary drivers of bacterial capsule maintenance in Klebsiella. 2021.

[6] Frederic D. Bushman, Kevin McCormick, and Scott Sherrill-Mix. Virus structures constrain transmission modes. *Nature Microbiology*, 4(11):1778–1780, July 2019.

[7] L.N. Calhoun and Y.M. Kwon. Structure, function and regulation of the DNA-binding protein Dps and its role in acid and oxidative stress resistance in Escherichia coli: a review: Escherichia coli Dps protein. *Journal of Applied Microbiology*, 110(2):375–386, February 2011.

[8] Spencer Cesar, Lisa Willis, and Kerwyn Casey Huang. Bacterial respiration during stationary phase induces intracellular damage that leads to delayed regrowth. *iScience*, 25(3):103765, March 2022.

[9] S. Chandrasekhar. Stochastic Problems in Physics and Astronomy. *Reviews of Modern Physics*, 15(1):1–89, January 1943.

[10] Emilia Chiancone and Pierpaolo Ceci. The multifaceted capacity of Dps proteins to combat bacterial stress conditions: Detoxification of iron and hydrogen peroxide and DNA binding. *Biochimica et Biophysica Acta (BBA) - General Subjects*, 1800(8):798–805, August 2010.

[11] Estelle E Clerc and et al. Strong chemotaxis by marine bacteria towards polysaccharides is enhanced by the abundant organosulfur compound DMSP. *Nature Communications*, 14:8080, 2023.

[12] Manoshi S. Datta, Elzbieta Sliwerska, Jeff Gore, Martin F. Polz, and Otto X. Cordero. Microbial interactions lead to rapid micro-scale successions on model marine particles. *Nature Communications*, 7:11965, June 2016.

[13] Jenny Davis and Ronald Benner. Quantitative estimates of labile and semi-labile dissolved organic carbon in the western Arctic Ocean: A molecular approach. *Limnol. Oceanogr*, 52(6)(6):2434–2444, 2007.

[14] Tim N. Enke, Manoshi S. Datta, Julia Schwartzman, Nathan Cermak, Désirée Schmitz, Julien Barrere, Alberto Pascual-García, and Otto X. Cordero. Modular Assembly of Polysaccharide-Degrading Marine Microbial Communities. *Current Biology*, 29(9):1528–1535.e6, May 2019.

[15] Benjamin Ezraty, Alexandra Gennaris, Frédéric Barras, and Jean-François Collet. Oxidative stress, protein damage and repair in bacteria. *Nature Reviews Microbiology*, 15(7):385–396, July 2017.

[16] Matteo P. Ferla and Wayne M. Patrick. Bacterial methionine biosynthesis. *Microbiology*, 160(8):1571–1584, August 2014.

[17] Jiunn C. N. Fong, Khalid A. Syed, Karl E. Klose, and Fitnat H. Yildiz. Role of Vibrio polysaccharide (vps) genes in VPS production, biofilm formation and Vibrio cholerae pathogenesis. *Microbiology*, 156(9):2757–2769, September 2010.

[18] Gill G. Geesey and Richard Y. Morita. Capture of Arginine at Low Concentrations by a Marine Psychrophilic Bacterium. *Applied and Environmental Microbiology*, 38(6):1092–1097, December 1979.

[19] D. L. Gibson, A. P. White, S. D. Snyder, S. Martin, C. Heiss, P. Azadi, M. Surette, and W. W. Kay. *Salmonella* Produces an O-Antigen Capsule Regulated by AgfD and Important for Environmental Persistence. *Journal of Bacteriology*, 188(22):7722–7730, November 2006.

[20] B S Goldman and J R Roth. Genetic structure and regulation of the cysG gene in Salmonella typhimurium. *Journal of Bacteriology*, 175(5):1457–1466, March 1993.

[21] Laura Gómez-Consarnau, Neelam Akram, Kristoffer Lindell, Anders Pedersen, Richard Neutze, Debra L. Milton, José M. González, and Jarone Pinhassi. Proteorhodopsin Phototrophy Promotes Survival of Marine Bacteria during Starvation. *PLoS Biology*, 8(4):e1000358, April 2010.

[22] Jye-Lin Hsu, Hsuan-Cheng Chen, Hwei-Ling Peng, and Hwan-You Chang. Characterization of the Histidine-containing Phosphotransfer Protein B-mediated Multistep Phosphorelay System in Pseudomonas aeruginosa PAO1. *Journal of Biological Chemistry*, 283(15):9933–9944, April 2008.

[23] Xin Huang, Thomas Nero, Ranjuna Weerasekera, Katherine H. Matej, Alex Hinbest, Zhaowei Jiang, Rebecca F. Lee, Longjun Wu, Cecilia Chak, Japinder Nijjer, Isabella Gibaldi, Hang Yang, Nathan Gamble, Wai-Leung Ng, Stacy A. Malaker, Kaelyn Sumigray, Rich Olson, and Jing Yan. Vibrio cholerae biofilms use modular adhesins with glycan-targeting and nonspecific surface binding domains for colonization. *Nature Communications*, 14(1):2104, April 2023.

[24] Dana E. Hunt, Lawrence A. David, Dirk Gevers, Sarah P. Preheim, Eric J. Alm, and Martin F. Polz. Resource Partitioning and Sympatric Differentiation Among Closely Related Bacterioplankton. *Science*, 320(5879):1081–1085, May 2008.

[25] Haiyan Huo, Rui He, Rongjing Zhang, and Junhua Yuan. Swimming *Escherichia coli* explore the environment by Lévy walk. *Applied and Environmental Microbiology*, 87(6):e02429–20, January 2021.

[26] James A. Imlay. The molecular mechanisms and physiological consequences of oxidative stress: lessons from a model bacterium. *Nature Reviews Microbiology*, 11(7):443–454, July 2013.

[27] James A. Imlay. Where in the world do bacteria experience oxidative stress?: Oxidative stress in natural environments. *Environmental Microbiology*, 21(2):521–530, February 2019.

[28] Keiichi Inoue, Hikaru Ono, Rei Abe-Yoshizumi, Susumu Yoshizawa, Hiroyasu Ito, Kazuhiro Kogure, and Hideki Kandori. A light-driven sodium ion pump in marine bacteria. *Nature Communications*, 4(1):1678, April 2013.

[29] Christoph Kaleta, Sascha Schäuble, Ursula Rinas, and Stefan Schuster. Metabolic costs of amino acid and protein production in *Escherichia coli*. *Biotechnology Journal*, 8(9):1105–1114, September 2013.

[30] Peter D Karp, Richard Billington, Ron Caspi, Carol A Fulcher, Mario Latendresse, Anamika Kothari, Ingrid M Keseler, Markus Krummenacker, Peter E Midford, Quang Ong, Wai Kit Ong, Suzanne M Paley, and Pallavi Subhraveti. The BioCyc collection of microbial genomes and metabolic pathways. *Briefings in Bioinformatics*, 20(4):1085–1093, July 2019.

[31] Rg Keil and Dl Kirchman. Utilization of dissolved protein and amino acids in the northern Sargasso Sea. *Aquatic Microbial Ecology*, 18:293–300, 1999.

[32] Thomas Kiørboe. *A mechanistic approach to plankton ecology*. Princeton University Press, 2008.

[33] Arne Klingner, Annekathrin Bartsch, Marco Dogs, Irene Wagner-Döbler, Dieter Jahn, Meinhard Simon, Thorsten Brinkhoff, Judith Becker, and Christoph Wittmann. Large-Scale $^{13}$ C Flux Profiling Reveals Conservation of the Entner-Doudoroff Pathway as a Glycolytic Strategy among Marine Bacteria That Use Glucose. *Applied and Environmental Microbiology*, 81(7):2408–2422, April 2015.

[34] Bennett S. Lambert, Vicente I. Fernandez, and Roman Stocker. Motility drives bacterial encounter with particles responsible for carbon export throughout the ocean. *Limnology and Oceanography Letters*, 4(5):113–118, October 2019.

[35] Ganhui Lan, Pablo Sartori, Silke Neumann, Victor Sourjik, and Yuhai Tu. The energy–speed–accuracy trade-off in sensory adaptation. *Nature Physics*, 8(5):422–428, May 2012.

[36] Marlon R. Lewis, David Hebert, W. Glen Harrison, Trevor Platt, and Neil S. Oakey. Vertical Nitrate Fluxes in the Oligotrophic Ocean. *Science*, 234(4778):870–873, November 1986.

[37] A Martinez and R Kolter. Protection of DNA during oxidative stress by the nonspecific DNA-binding protein Dps. *Journal of Bacteriology*, 179(16):5188–5194, August 1997.

[38] Miguel A Matilla and Tino Krell. The effect of bacterial chemotaxis on host infection and pathogenicity. *FEMS Microbiology Reviews*, 42(1), January 2018.

[39] Diane McDougald, Lan Gong, Sujatha Srinivasan, Erika Hild, Lyndal Thompson, Scott A Rice, and S Kjelleberg. Defences against oxidative stress during starvation in bacteria. *Van Leeuwenhoek*, 2002.

[40] Bui Quang Minh, Heiko A Schmidt, Olga Chernomor, Dominik Schrempf, Michael D Woodhams, Arndt Von Haeseler, and Robert Lanfear. IQ-TREE 2: New Models and Efficient Methods for Phylogenetic Inference in the Genomic Era. *Molecular Biology and Evolution*, 37(5):1530–1534, May 2020.

[41] J. Jeffrey Morris, Zackary I. Johnson, Martin J. Szul, Martin Keller, and Erik R. Zinser. Dependence of the Cyanobacterium Prochlorococcus on Hydrogen Peroxide Scavenging Microbes for Growth at the Ocean's Surface. *PLoS ONE*, 6(2):e16805, February 2011.

[42] J. Jeffrey Morris, Richard E. Lenski, and Erik R. Zinser. The Black Queen Hypothesis: Evolution of Dependencies through Adaptive Gene Loss. *mBio*, 3(2):e00036–12, May 2012.

[43] J. Jeffrey Morris, Andrew L. Rose, and Zhiying Lu. Reactive oxygen species in the world ocean and their impacts on marine ecosystems. *Redox Biology*, 52:102285, June 2022.

[44] Noele Norris, Naomi M. Levine, Vicente I. Fernandez, and Roman Stocker. Mechanistic model of nutrient uptake explains dichotomy between marine oligotrophic and copiotrophic bacteria. *PLOS Computational Biology*, 17(5):e1009023, May 2021.

[45] Martin Ostrowski, Ricardo Cavicchioli, Maarten Blaauw, and Jan C. Gottschal. Specific Growth Rate Plays a Critical Role in Hydrogen Peroxide Resistance of the Marine Oligotrophic Ultramicrobacterium *Sphingomonas alaskensis* Strain RB2256. *Applied and Environmental Microbiology*, 67(3):1292–1299, March 2001.

[46] Mark Pagel. Detecting Correlated Evolution on Phylogenies: A General Method for the Comparative Analysis of Discrete Characters. *Proceedings: Biological Sciences*, 255(1342):37–45, 1994.

[47] Marc Picheral, Sarah Searson, V. Taillandier, Annick Bricaud, Emmanuel Boss, Lars Stemmann, G. Gorsky, Coordinators Tara Oceans Consortium, and Participants Tara Oceans Expedition. Vertical profiles of environmental parameters measured from physical, optical and imaging sensors during Tara Oceans expedition 2009-2013, 2014. Type: data set.

[48] Jean-Baptiste Raina, Vicente Fernandez, Bennett Lambert, Roman Stocker, and Justin R. Seymour. The role of microbial motility and chemotaxis in symbiosis. *Nature Reviews Microbiology*, 17(5):284–294, May 2019.

[49] Christopher V. Rao, George D. Glekas, and George W. Ordal. The three adaptation systems of Bacillus subtilis chemotaxis. *Trends in Microbiology*, 16(10):480–487, October 2008.

[50] Liam J. Revell. phytools 2.0: an updated R ecosystem for phylogenetic comparative methods (and other things). *PeerJ*, 12:e16505, January 2024.

[51] Abraham. Savitzky and M. J. E. Golay. Smoothing and Differentiation of Data by Simplified Least Squares Procedures. *Analytical Chemistry*, 36(8):1627–1639, July 1964.

[52] D Schellenberg and E Furlongs. Resolution of the Multiplicity of the Glutamate and Aspartate Transport Systems of Escherichia coli. *Journal of Biological Chemistry*, 252(24):9055–9064, 1977.

[53] Severin J. Schink, Elena Biselli, Constantin Ammar, and Ulrich Gerland. Death Rate of E. coli during Starvation Is Set by Maintenance Cost and Biomass Recycling. *Cell Systems*, 9(1):64–73.e3, July 2019.

[54] Carmen Schwechheimer, Kassidy Hebert, Sarvind Tripathi, Praveen K. Singh, Kyle A. Floyd, Elise R. Brown, Monique E. Porcella, Jacqueline Osorio, Joseph T. M. Kiblen, Fernando A. Pagliai, Knut Drescher, Seth M. Rubin, and Fitnat H. Yildiz. A tyrosine phosphoregulatory system controls exopolysaccharide biosynthesis and biofilm formation in Vibrio cholerae. *PLOS Pathogens*, 16(8):e1008745, August 2020.

[55] T. S. Shimizu, N. Delalez, K. Pichler, and H. C. Berg. Monitoring bacterial chemotaxis by using bioluminescence resonance energy transfer: Absence of feedback from the flagellar motors. *Proceedings of the National Academy of Sciences*, 103(7):2093–2097, February 2006.

[56] William R. Shoemaker, Stuart E. Jones, Mario E. Muscarella, Megan G. Behringer, Brent K. Lehmkuhl, and Jay T. Lennon. Microbial population dynamics and evolutionary outcomes under extreme energy limitation. *Proceedings of the National Academy of Sciences*, 118(33):e2101691118, August 2021.

[57] Kwangmin Son, Filippo Menolascina, and Roman Stocker. Speed-dependent chemotactic precision in marine bacteria. *Proceedings of the National Academy of Sciences*, 113(31):8624–8629, August 2016.

[58] Vanessa Sperandio, Alfredo G. Torres, and James B. Kaper. Quorum sensing *Escherichia coli* regulators B and C (QseBC): a novel two-component regulatory system involved in the regulation of flagella and motility by quorum sensing in *E. coli*: QseBC regulates flagella and motility in *E. coli*. *Molecular Microbiology*, 43(3):809–821, February 2002.

[59] Laura Steindler, Michael S. Schwalbach, Daniel P. Smith, Francis Chan, and Stephen J. Giovannoni. Energy Starved Candidatus Pelagibacter Ubique Substitutes Light-Mediated ATP Production for Endogenous Carbon Respiration. *PLoS ONE*, 6(5):e19725, May 2011.

[60] Ke Stoderegger and Gj Herndl. Production of exopolymer particles by marine bacterioplankton under contrasting turbulence conditions. *Marine Ecology Progress Series*, 189:9–16, 1999.

[61] Johannes Taktikos, Holger Stark, and Vasily Zaburdaev. How the Motility Pattern of Bacteria Affects Their Dispersal and Chemotaxis. *PLoS ONE*, 8(12):e81936, December 2013.

[62] Baishnab C Tripathy, Irena Sherameti, and Ralf Oelmüller. Siroheme: An essential component for life on earth. *Plant Signaling & Behavior*, 5(1):14–20, January 2010.

[63] Lam Si Tung Ho and Cécile Ané. A Linear-Time Algorithm for Gaussian and Non-Gaussian Trait Evolution Models. *Systematic Biology*, 63(3):397–408, May 2014.

[64] G. M. Viswanathan, Sergey V. Buldyrev, Shlomo Havlin, M. G. E. da Luz, E. P. Raposo, and H. Eugene Stanley. Optimizing the success of random searches. *Nature*, 401(6756):911–914, October 1999.

[65] J. M. Walter, D. Greenfield, C. Bustamante, and J. Liphardt. Light-powering Escherichia coli with proteorhodopsin. *Proceedings of the National Academy of Sciences*, 104(7):2408–2412, February 2007.

[66] Nicholas J. Watmough and Frank E. Frerman. The electron transfer flavoprotein: Ubiquinone oxidoreductases. *Biochimica et Biophysica Acta (BBA) - Bioenergetics*, 1797(12):1910–1916, December 2010.

[67] Ben-Haim Y. and Rosenberg E. A novel Vibrio sp. pathogen of the coral Pocillopora damicornis. *Marine Biology*, 141(1):47–55, July 2002.

[68] Fitnat H. Yildiz and Karen L. Visick. Vibrio biofilms: so much the same yet so different. *Trends in Microbiology*, 17(3):109–118, March 2009.

[69] Mattia Zampieri, Manuel Hörl, Florian Hotz, Nicola F. Müller, and Uwe Sauer. Regulatory mechanisms underlying coordination of amino acid and glucose catabolism in Escherichia coli. *Nature Communications*, 10(1):3354, December 2019.

[70] Yucheng Zheng, Huan Wang, Limin Huang, Tongchao Zhang, Bingbing Zong, Xuanxiu Ren, Yongwei Zhu, Fangyu Song, Xiangru Wang, Huanchun Chen, and Chen Tan. Effect of O antigen ligase gene mutation on oxidative stress resistance and pathogenicity of NMEC strain RS218. *Microbial Pathogenesis*, 136:103656, November 2019.
